# TDP-43 dysregulation of polyadenylation site selection is a defining feature of RNA misprocessing in ALS/FTD and related disorders

**DOI:** 10.1101/2024.01.22.576709

**Authors:** Frederick J. Arnold, Ya Cui, Sebastian Michels, Michael R. Colwin, Cameron Stockford, Wenbin Ye, Oliver H. Tam, Sneha Menon, Wendy G. Situ, Kean C. K. Ehsani, Sierra Howard, Molly Gale Hammell, Wei Li, Albert R. La Spada

## Abstract

Nuclear clearance and cytoplasmic aggregation of the RNA-binding protein TDP-43 are observed in many neurodegenerative disorders, including amyotrophic lateral sclerosis (ALS) and fronto- temporal dementia (FTD). Although TDP-43 dysregulation of splicing has emerged as a key event in these diseases, TDP-43 can also regulate polyadenylation; yet, this has not been adequately studied. Here, we applied the dynamic analysis of polyadenylation from RNA-seq (DaPars) tool to ALS/FTD transcriptome datasets, and report extensive alternative polyadenylation (APA) upon TDP-43 alteration in ALS/FTD cell models and postmortem ALS/FTD neuronal nuclei. Importantly, many identified APA genes highlight pathways implicated in ALS/FTD pathogenesis. To determine the functional significance of APA elicited by TDP-43 nuclear depletion, we examined microtubule affinity regulating kinase 3 (MARK3). Nuclear loss of TDP-43 yielded increased expression of MARK3 transcripts with longer 3’UTRs, resulting in greater transcript stability and elevated MARK3 protein levels, which promotes increased neuronal tau S262 phosphorylation. Our findings define changes in polyadenylation site selection as a previously unrecognized feature of TDP-43-driven disease pathology in ALS/FTD and highlight a potentially novel mechanistic link between TDP-43 dysfunction and tau regulation.

---

While the underlying etiology of many age-related neurodegenerative disorders remains unknown, distinct neuronal subtypes and specific brain regions exhibit a common pathological hallmark of nuclear clearance and cytoplasmic aggregation of TAR DNA-binding protein 43 (TDP- 43) ^1^. TDP-43 pathology is present in motor neurons in ∼97% of patients with amyotrophic lateral sclerosis (ALS) and in the neocortex of ∼45% of individuals suffering from frontotemporal dementia (FTD). Furthermore, as many as 57% of Alzheimer’s disease (AD) patients display TDP-43 pathology, which correlates with more rapid disease progression and worse cognitive impairment ^2,3^. Cytoplasmic aggregates of TDP-43 have also been observed in Huntington’s disease ^4^ and chronic traumatic encephalopathy ^5^, and are emerging as the defining pathology for limbic-predominant age-related TDP-43 encephalopathy (LATE), a recently described dementia characterized by atrophy in the medial temporal lobes and frontal cortex, though not exclusive to this brain region ^6^. Consequently, heterogenous neurodegenerative disorders characterized by TDP-43 pathology may collectively be referred to as ‘TDP-43 proteinopathies‘ ^1^.

Under physiological conditions, TDP-43 is a predominantly nuclear RNA-binding protein (RBP) that directly binds over 6,000 RNAs ^7^. Prior studies of ALS/FTD have identified differentially expressed and alternatively spliced genes associated with TDP-43 nuclear loss of function. This includes a motor neuron growth and repair factor, stathmin-2 (STMN2), in which TDP-43 loss of nuclear function results in de-repression of a cryptic 3’ splice site in the *STMN2* gene, favoring inclusion of a cryptic exon in the *STMN2* pre-mRNA, premature transcription termination, and loss of STMN2 protein expression ^8–10^. Another gene subject to TDP-43 splicing dysregulation in ALS/FTD is *UNC13A*, as depletion of TDP-43 in the nucleus promotes cryptic exon inclusion within the *UNC13A* pre-mRNA, resulting in loss of UNC13A protein expression ^11,12^. Both STMN2 and UNC13A are currently under investigation as candidates for therapeutic intervention and biomarker development in ALS/FTD.

In addition to its role in RNA splicing, TDP-43 regulates alternative polyadenylation (APA) by binding target RNAs near polyadenylation signals (PAS) ^13^, yet this aspect of TDP-43 function remains relatively unexplored in TDP-43 proteinopathies. Notably, up to 70% of all human pre-mRNAs contain more than one polyadenylation (poly(A)) site, and differential PAS utilization yields transcripts with distinct 3’ untranslated regions (3’UTRs) ^14,15^. APA is a highly conserved and ubiquitous mechanism of gene regulation that is exceedingly cell-type specific ^16^. As it turns out, neurons tend to preferentially utilize distal poly(A) sites ^17,18^, accounting for the well-established observation that genes expressed in the central nervous system (CNS) encode transcripts containing the longest 3’UTR sequences of all tissues throughout the body. This suggests that APA is likely an important post-transcriptional regulatory mechanism in neurons and other CNS cell types, as the 3’UTR is highly enriched for binding motifs for microRNAs (miRNAs) ^19^ and RBPs ^20^. Hence, by modulating the presence or absence of such cis-regulatory elements, APA can produce transcript isoforms with substantial differences in RNA stability, subcellular localization, nuclear export, and transcription efficiency ^21^. These changes in RNA metabolism can markedly impact protein expression, though it is also likely that changes in subcellular localization can modulate gene function without affecting steady-state RNA levels. An important consideration for the study of APA is that standard informatics analysis of RNA-seq data does not fully capture APA events. For this reason, a growing number of computational methods have been developed to infer PAS usage from standard, short-read, bulk RNA-seq transcriptome data, allowing post-hoc analysis of APA from existing datasets ^22^.

Here we applied the dynamic analysis of polyadenylation from RNA-sequencing (DaPars) tool to transcriptome data sets obtained from neuron cell culture models of ALS/FTD and from ALS/FTD patient postmortem neuron nuclei. We identified hundreds of previously unknown genes regulated by TDP-43, with many genes conserved across multiple cell types and model systems. As a substantial number of these genes function in cellular pathways implicated in ALS/FTD disease pathogenesis, APA analysis can provide insight into how TDP-43 dysfunction alters the transcriptome of neurons and other cell types. To determine the biological significance of TDP- 43 APA, we studied its effects on the kinase MARK3, a highly significant hit from APA analysis of ALS/FTD frontal cortex neuron nuclei containing or depleted of TDP-43. We confirmed that upon TDP-43 knock-down, the *MARK3* gene displayed increased RNA stability resulting in elevated MARK3 protein expression in iPSC-derived motor neurons, and we documented that TDP-43 depletion promoted increased phosphorylation of tau at serine 262, a previously established read-out for tau dysfunction in Alzheimer’s disease ^23^. Our results indicate that TDP-43 dysregulation of polyadenylation site selection is another facet of the RNA processing dysfunction which is a central feature of the disease process in TDP-43 proteinopathies.

## Results

### TDP-43 depletion induces widespread alternative polyadenylation in neuronal cells

To identify alternative polyadenylation (APA) events regulated by TDP-43 in neuronal cells, we employed the dynamic analysis of polyadenylation from RNA-sequencing (DaPars) tool to previously published RNA-sequencing datasets (**Fig. 1a**). While a number of tools have been developed to quantify APA from bulk RNA-seq data ^22^, we selected DaPars ^24^, which identifies and quantifies *de novo* polyA site usage by applying a linear regression model to localized changes in RNA-seq read density within the 3’UTR. Given the established role of TDP-43 in regulating the splicing of unannotated ‘cryptic’ exons, we hypothesized that ‘cryptic,’ unannotated polyA sites may be similarly utilized upon TDP-43 depletion or mutation. DaPars calculates the percentage of distal polyA site usage index (PDUI) for each transcript, with the results averaged across replicates to determine ΔPDUI in cells where TDP-43 was knocked down or mutated in comparison to control conditions. A positive ΔPDUI denotes increased relative expression of transcripts with a longer 3’UTR, while a negative ΔPDUI signifies increased relative expression of transcripts with a shorter 3’UTR.

**Figure 1.**
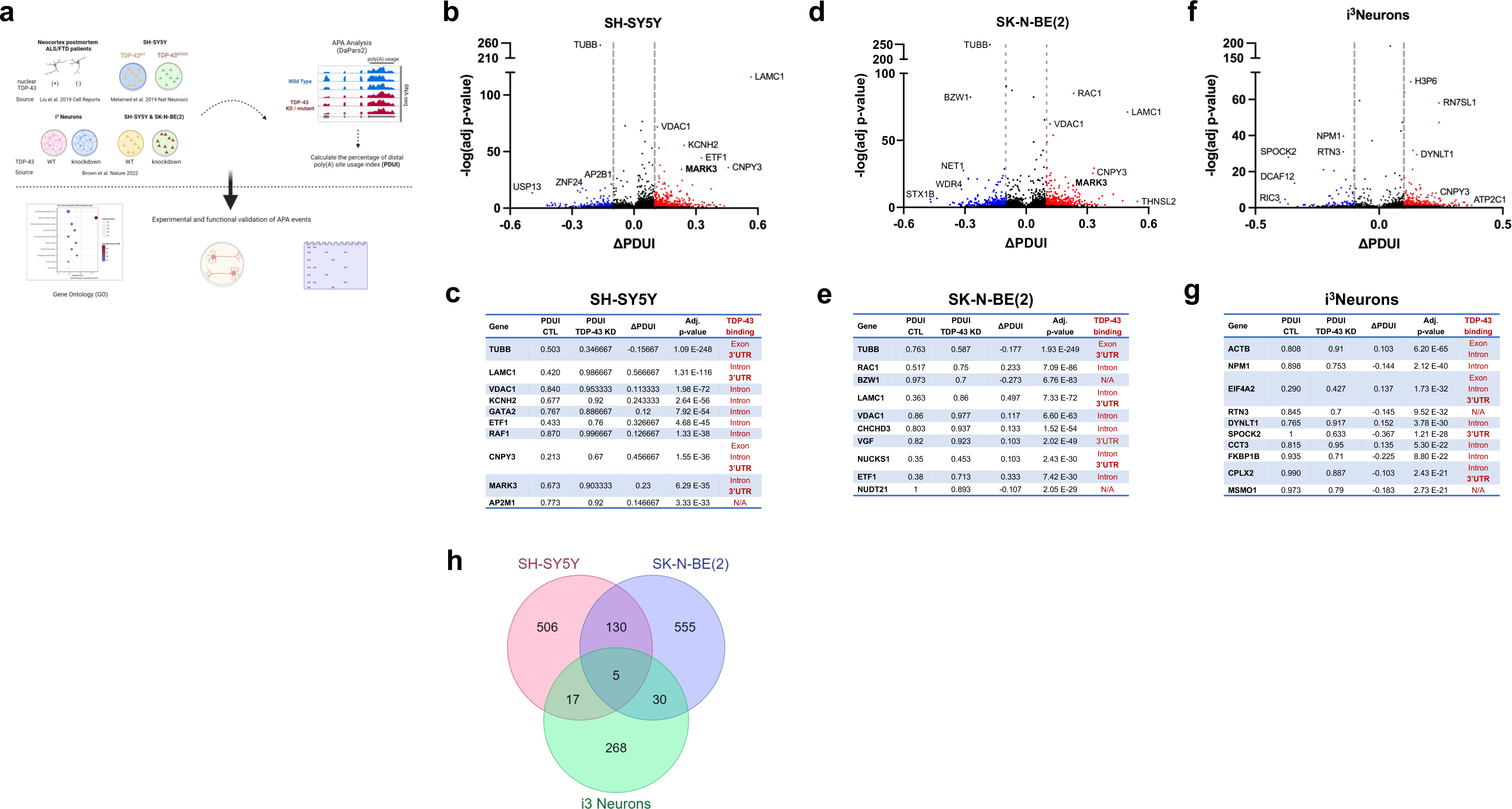
TDP-43 depletion induces widespread alternative polyadenylation. (**a**) Overview of the study. We applied the dynamic analysis of polyadenylation from RNA-seq (DaPars) tool to published RNA-seq datasets to detect APA events resulting from TDP-43 depletion or mutation in immortalized cell lines, stem cell-derived neurons, and postmortem ALS/FTD neocortex. We then performed experimental and functional validation of select targets. (**b, d, f**) Volcano plots depicting APA events in SH-SY5Y cells (**b**), SK-N-BE(2) cells (**d**), and i^3^Neurons (**f**) in which TDP-43 was knocked down via shRNA treatment. APA genes with FDR adj. p value < 0.05 and ΔPDUI ý 0.1 are depicted in red, and APA genes with FDR adj. p value < 0.05 and ΔPDUI ý 0.1 are depicted in blue. (**c, e, g**) Table of top 10 APA events in SH- SY5Y cells (**c**), SK-N-BE(2) cells (**e**), and i^3^Neurons (**g**) by FDR adj. p-value with ƖΔPDUIƖ ≥ 0.1, along with evidence of TDP-43 binding sites in each gene by eCLIP-seq (red). (**h**) Venn diagram illustrating the intersection of APA events across datasets.

Using published RNA-seq data from SH-SY5Y cells, SK-N-BE(2) cells, and i^3^Neurons in which TDP- 43 was depleted by shRNA knock-down ^11^, we identified hundreds of genes in which TDP-43 knock-down resulted in significant APA (FDR adj p < 0.05, |ΔPDUI| ý 0.1) (**Fig. 1b-g, Supplementary Tables 1-3**). APA genes exhibit both 3’UTR lengthening and shortening upon TDP-43 knock-down; however, we observed increased use of distal polyA sites in 69.9% (460/658) of APA genes in SH-SY5Y cells, 59.6% (429/720) of APA genes in SK-N-BE(2) cells, and 78.1% (250/320) of APA genes in i^3^Neurons. Nearly 20% of APA events were shared between the closely related SH-SY5Y and SK-N-BE(2) neuroblastoma cell lines (**Fig. 1h**), including 4/10 of the top APA genes by p value (**Fig. 1c, e**). Additionally, 16.3% (52/320) of APA genes in i^3^Neurons overlapped with at least one of the immortalized neuronal cell lines (**Fig. 1h**), indicating that, while APA is known to be highly cell-type specific ^16^, many APA events regulated by TDP-43 are conserved across multiple neuronal cell types.

We also applied DaPars to previously published transcriptome data sets in which TDP-43 was depleted by siRNAs in SH-SY5Y cells ^10^ or in human embryonic stem cell derived motor neurons (hESC-MNs) for 96 hrs ^8^ (**Supplementary Fig. 1, Supplementary Tables 4, 5**). As expected, we observed fewer APA changes in cells in which TDP-43 was knocked down for a shorter duration or incompletely (∼60% in hESC-MNs). Both datasets do, however, exhibit significantly decreased PDUI in *SMC1A*, as previously shown for HEK293 cells upon TDP-43 siRNA knock-down ^13^.

Given that APA can affect cellular function by regulating mRNA stability and subcellular localization ^21^, we performed gene ontology (GO) analysis of APA genes in each neuronal cell type (**Supplementary** Fig. 2). Shared terms between datasets for biological pathways known to be highly relevant to ALS/FTD included: ‘establishment of protein localization’ (SH-SY5Y and i^3^Neurons) and ‘cytoskeleton organization’ (SH-SY5Y and SK-N-BE(2)), and shared terms for GO molecular functions were: ‘kinase activity’, ‘protein kinase binding’, and ‘cytoskeletal protein binding’ (SH-SY5Y and SK-N-BE(2)). These results indicate that TDP-43 regulation of APA impacts disease-relevant pathways and should be considered, along with differential expression and alternative splicing, as a key aspect of TDP-43 dysfunction.

### APA genes directly bound by TDP-43 in the 3’UTR exhibit preferential 3’UTR-lengthening upon TDP-43 depletion

By cross-referencing significant APA genes with TDP-43 eCLIP-seq data from SH-SY5Y cells ^25^, we found that TDP-43 directly binds either within the 3’UTR or in another region of the transcript to 43.2% (284/658) of APA genes in SH-SY5Y cells, 47.4% (341/720) of APA genes in SK-N-BE(2) cells, and 51.3% (164/320) of APA genes in i^3^Neurons, (**Fig. 2a**, **Supplementary** Fig. 3). In neuronal cells chronically depleted of TDP-43, it is likely that cytotoxicity contributes to gene regulatory changes that do not necessarily reflect direct regulation by TDP-43. Indeed, we noted that in i^3^Neurons, only 36.7% of differentially expressed genes are targets of TDP-43 binding by eCLIP-seq, while a significantly higher proportion of cryptic exon (60.3%) and APA (51.3%) events are observed in genes directly bound by TDP-43 (**Fig. 2b**). In support of our overall finding that TDP-43 depletion preferentially increases use of a distal PAS, we found that APA genes exhibit increased PDUI upon TDP-43 loss – when TDP-43 binds within the 3’UTR, with i^3^Neurons displaying the most significant trend (**Fig. 2c-e**; see also accompanying paper from Gitler and colleagues ^26^).

**Figure 2.**
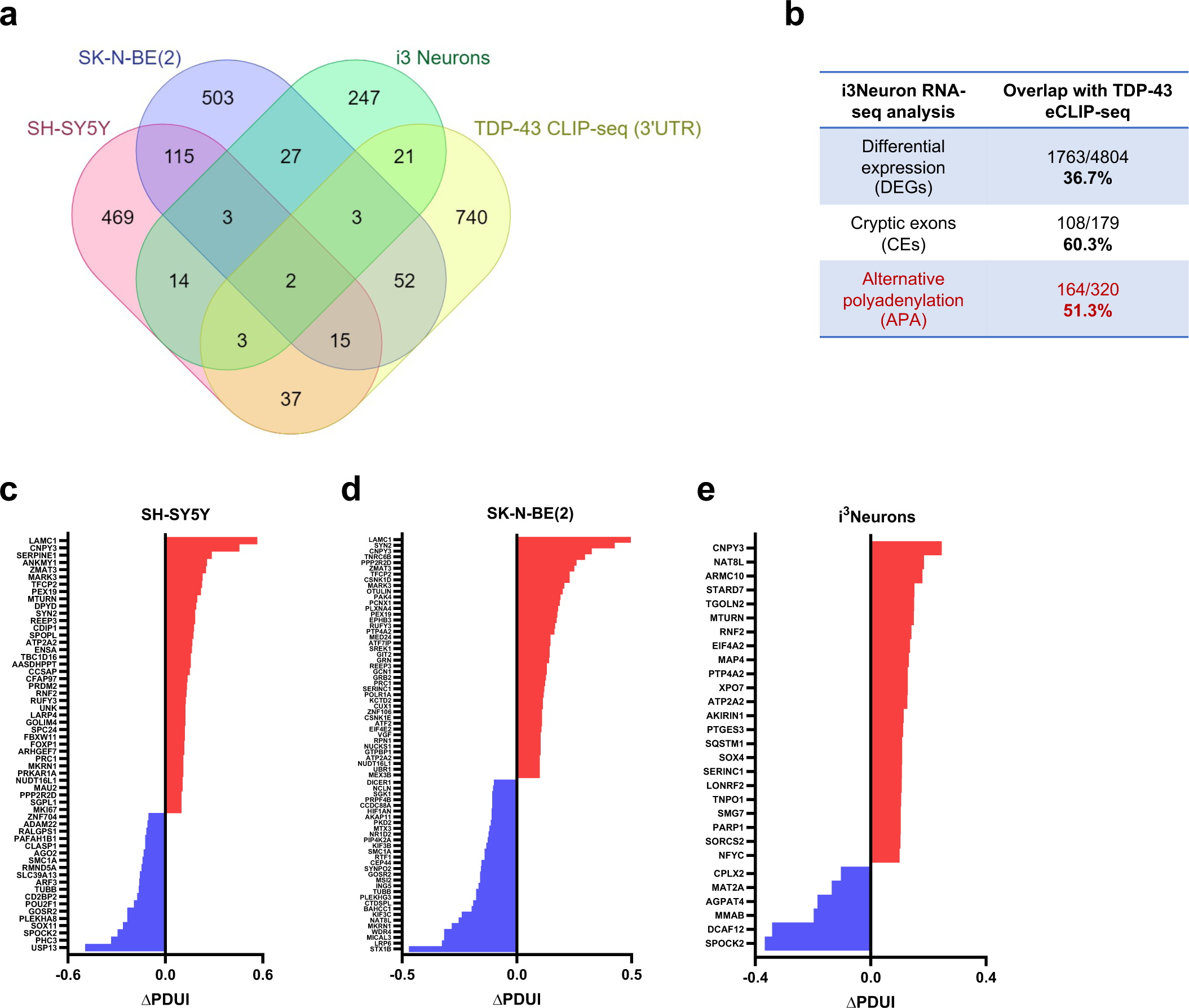
APA genes directly bound by TDP-43 in the 3’UTR exhibit preferential 3’UTR- lengthening upon TDP-43 depletion. (**a**) Venn diagram illustrating the proportion of APA genes for which there is published evidence of TDP-43 binding within the 3’UTR. (**b**) Percentage of differentially expressed genes (DEGs), cryptic exon (CE)-containing genes, and APA genes that contain TDP-43 binding sites from RNA- seq of i^3^Neurons +/- TDP-43 knock-down. (**c, d, e**) Graphical representation of ΔPDUI for all APA genes with TDP-43 binding sites in the 3’UTR.

### Experimental and functional validation of TDP-43 APA genes in neuronal cells

To experimentally validate TDP-43 APA events, we first utilized the above SH-SY5Y cell model, in which a doxycycline-inducible shRNA against TDP-43 is stably integrated ^11^. We selected two APA genes to validate based on their robust and consistent shift in PDUI upon TDP-43 knockdown, and because TDP-43 binds directly within each 3’UTR (**Supplementary** Fig. 4). The Canopy FGF signaling regulator 3 (*CNPY3*) gene was among the 3 greatest PDUI increases in each of our tested neuronal cell lines (**Fig. 2c-e**). *CNPY3* exhibits extensive alternative splicing, with numerous annotated transcript variants. In examining the RNA-seq tracks for *CNPY3*, we found that TDP-43 knock-down significantly increases PAS usage within a specific transcript variant (NM_001318848.2, ‘variant two’), defined by an alternative last exon without an upstream splice junction. In line with this data, we confirmed by RT-PCR that TDP-43 depletion results in a significant increase in the use of the variant two 3’UTR relative to variant one (NM_006586.5) (**Fig. 3a**). *CNPY3* APA was similarly described in the accompanying manuscript by Bryce-Smith et al. ^27^, and was independently detected by Zeng et al. in i^3^Neurons depleted of TDP-43 in their accompanying study ^26^. Increased production of an alternative *CNPY3* isoform upon TDP-43 depletion similarly parallels findings from Zeng et al. in which TDP-43 knock-down resulted in increased expression of an alternative *SFPQ* isoform ^26^.

**Figure 3.**
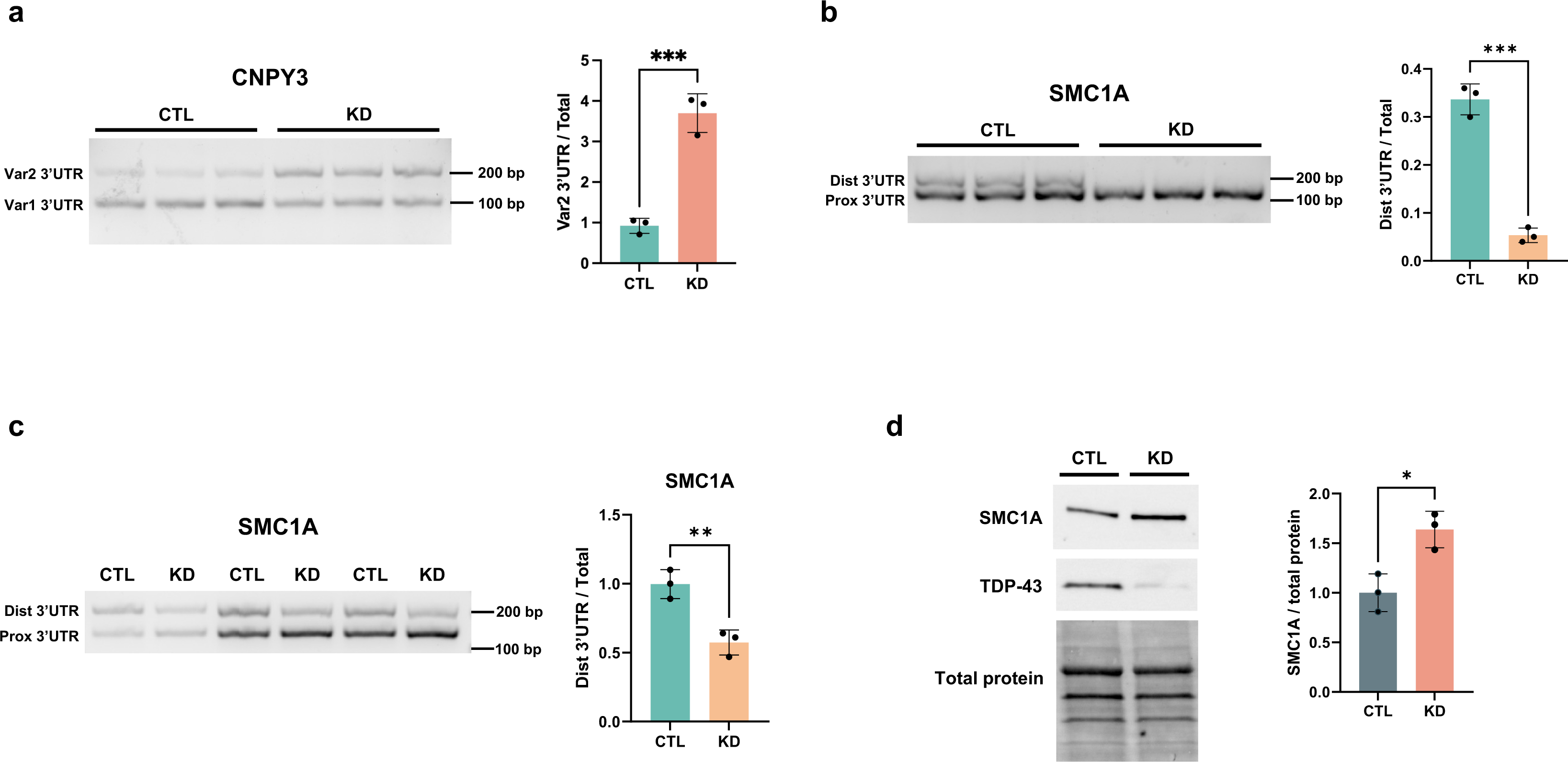
Experimental and functional validation of TDP-43 APA genes in neuronal cells. (**a**) PCR validation of increased usage of the *CNPY3* variant two 3’UTR upon TDP-43 knock- down in SH-SY5Y cells. (**b**) PCR validation of increased proximal PAS usage of SMC1A upon TDP-43 knock-down in SH-SY5Y cells. (**c**) Validation of SMC1A APA in iPSC-derived motor neurons (iPSC-MNs) in which TDP-43 was knocked down with shRNA for 10 days. (**d**) Immunoblot analysis of SMC1A in iPSC-MNs reveals that *SMC1A* APA corresponds with an increase in SMC1A protein levels. *p<0.05, **p<0.01, ***p<0.001; two-tailed t-test; n = 3 biological replicates. Data are presented as mean values ± SEM.

In contrast with *CNPY3*, DaPars calculated a significant decrease in distal PAS usage for *SMC1A* (**Supplementary** Fig. 4b), which we experimentally validated by RT-PCR in SH-SY5Y cells (**Fig. 3b**). As noted, APA of *SMC1A* was previously characterized in HEK293 cells upon TDP-43 knockdown, indicating that this is a highly sensitive APA event across diverse cell types. Given that SMC1A is a subunit of the cohesin complex and plays a critical role in chromatin organization, we sought to investigate the biological significance of SMC1A in neuronal cells. Using DIV38 induced pluripotent stem cell-derived motor neurons (iPSC-MNs) in which TDP-43 was knocked down for 10 days (**Supplementary** Fig. 5), we confirmed that TDP-43 loss indeed reduces use of the distal *SMC1A* PAS in human motor neurons (**Fig. 3c**). We further found that APA of *SMC1A* corresponds with a significant increase in SMC1A protein levels in iPSC-MNs (**Fig. 3d**), suggesting that altered regulation of *SMC1A* APA may contribute to impaired chromatin organization upon TDP-43 loss, as previously reported in postmortem neuronal nuclei ^28^.

### Mutant TDP-43 induces APA in genes that function in the oxidative stress response

To explore APA dysregulation that reflects TDP-43 gain-of-function in addition to loss-of-function pathology, we next applied DaPars to RNA-seq data generated from SH-SY5Y cells with homozygous mutation of TDP-43^N352S^ achieved via CRISPR-Cas9 genome editing ^10^. Consistent with prior studies, 72% (59/82) of significant APA events corresponded with increased use of a distal PAS (**Fig. 4a, Supplementary Table 6**). Notably, GO analysis revealed significant enrichment of genes that function in the ‘response to oxidative stress’ pathway (**Fig. 4b**). Numerous studies have implicated oxidative stress in ALS/FTD pathogenesis, including recent evidence that TDP-43 aggregation induces the generation of reactive oxygen species (ROS) ^29^. To determine if TDP-43^N352S^ SH-SY5Y cells display an altered oxidative stress response, we measured ROS at 30 minutes and 120 minutes after treating wildtype or TDP-43^N352S^ SH-SY5Y cells with hydrogen peroxide. While there was no initial difference in ROS levels between control and TDP- 43^N352^ cells, ROS levels were significantly higher in TDP-43^N352S^ SH-SY5Y cells compared to wildtype cells after 120 minutes of exposure (**Fig. 4c**). This provides proof-of-concept that TDP-43 APA genes function in cellular pathways implicated in neurodegenerative disease, and that characterization of APA events can highlight disease-relevant phenotypes in cell models of ALS.

**Figure 4.**
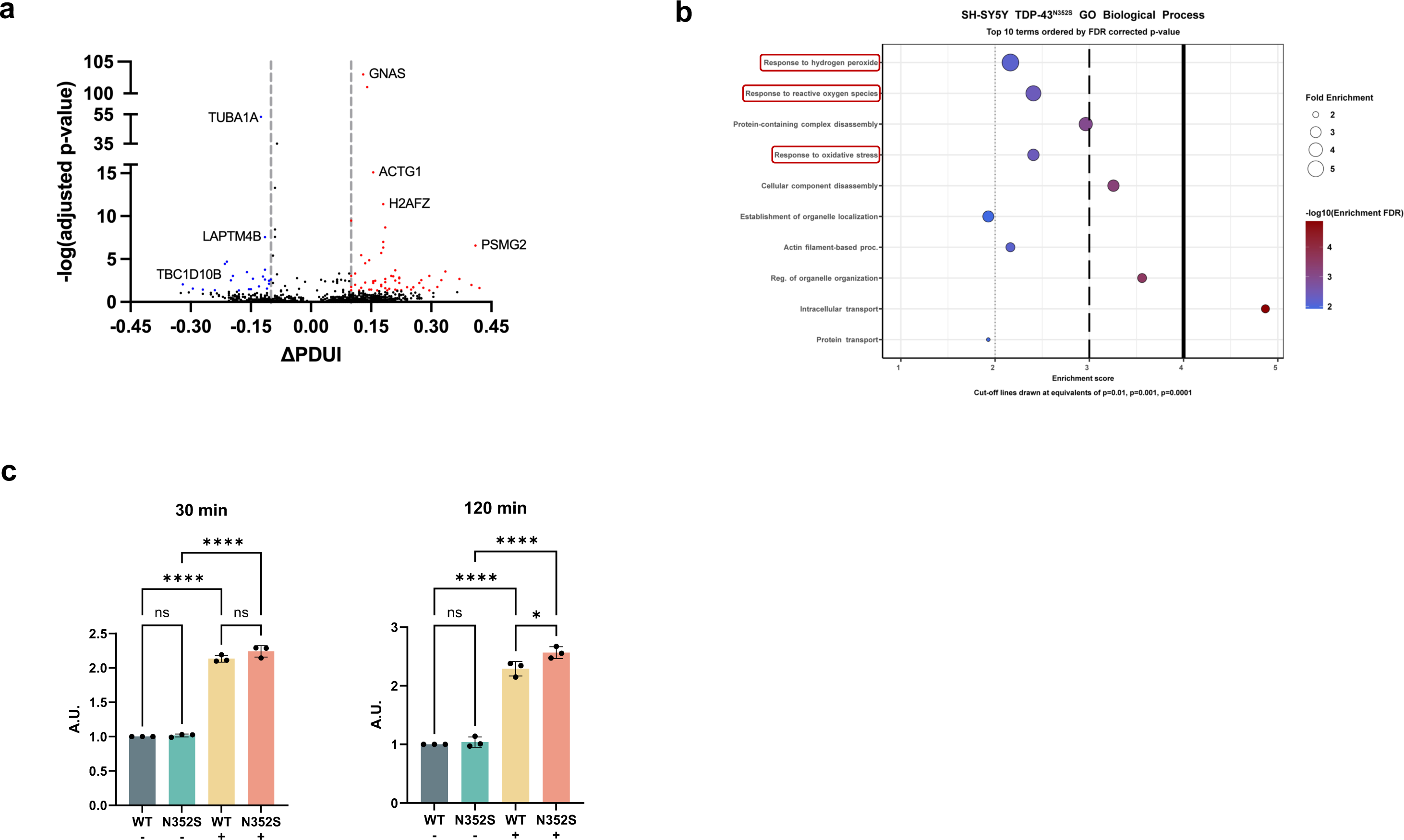
Mutant TDP-43 induces APA in genes functioning in the oxidative stress response. (**a**) Volcano plot depicting APA genes in SH-SY5Y cells expressing homozygous TDP-43^N352S^ via CRISPR-Cas9 mediated genome editing. APA genes with FDR adj. p value < 0.05 and ΔPDUI ý 0.1 are depicted in red, and APA genes with FDR adj. p value < 0.05 and ΔPDUI :: 0.1 are depicted in blue. (**b**) GO BP analysis of APA events in TDP-43^N352S^ cells reveals impaired response to hydrogen peroxide (H2O2). (**c**) Wildtype and TDP-43^N352S^ cells were treated with 100 μM H2O2. ROS was detected at 30 minutes and 120 minutes using a fluorometric intracellular ROS detection kit. *p<0.05. ****p<0.0001; one-way ANOVA; n = 3 biological replicates. Data are presented as mean values ± SEM.

### Nuclear clearance of TDP-43 induces APA in ALS/FTD patient neurons

To further determine the significance of TDP-43 APA dysregulation in human patients, we considered APA events in neuron nuclei obtained from seven post-mortem ALS/FTD neocortex samples, where FACS sorting resulted in transcriptome data sets for neurons either containing or depleted of nuclear TDP-43 ^28^. We identified 181 APA genes (|ΔPDUI| > 0.1, p<0.05) in neuronal nuclei lacking nuclear TDP-43 (**Fig. 5a, b, Supplementary Table 7**), but unlike in neuronal cell culture models, we observed a preference towards negative ΔPDUI in postmortem TDP-43 nuclear-depleted neurons, as 72.7% (63/87) of genes exhibited 3’UTR shortening. By correlating the most significant APA events (|ΔPDUI| > 0.1, FDR p<0.05) with eCLIP-seq of TDP- 43, we noted that 57.7% of these APA genes are bound by TDP-43 (**Fig. 5c**), suggesting that many APA events are directly regulated by TDP-43. We then performed GO analysis for APA events in coding mRNA with p < 0.05 and [|ΔPDUI| ≥ 0.1] and found significant enrichment of genes functioning in the ‘histamine response’, ‘synapse assembly’, and ‘protein transport’ pathways (**Fig. 5d**). These pathways have been previously implicated in ALS disease models, again underscoring potential contributions of APA events to ALS pathobiology ^30–32^.

**Figure 5.**
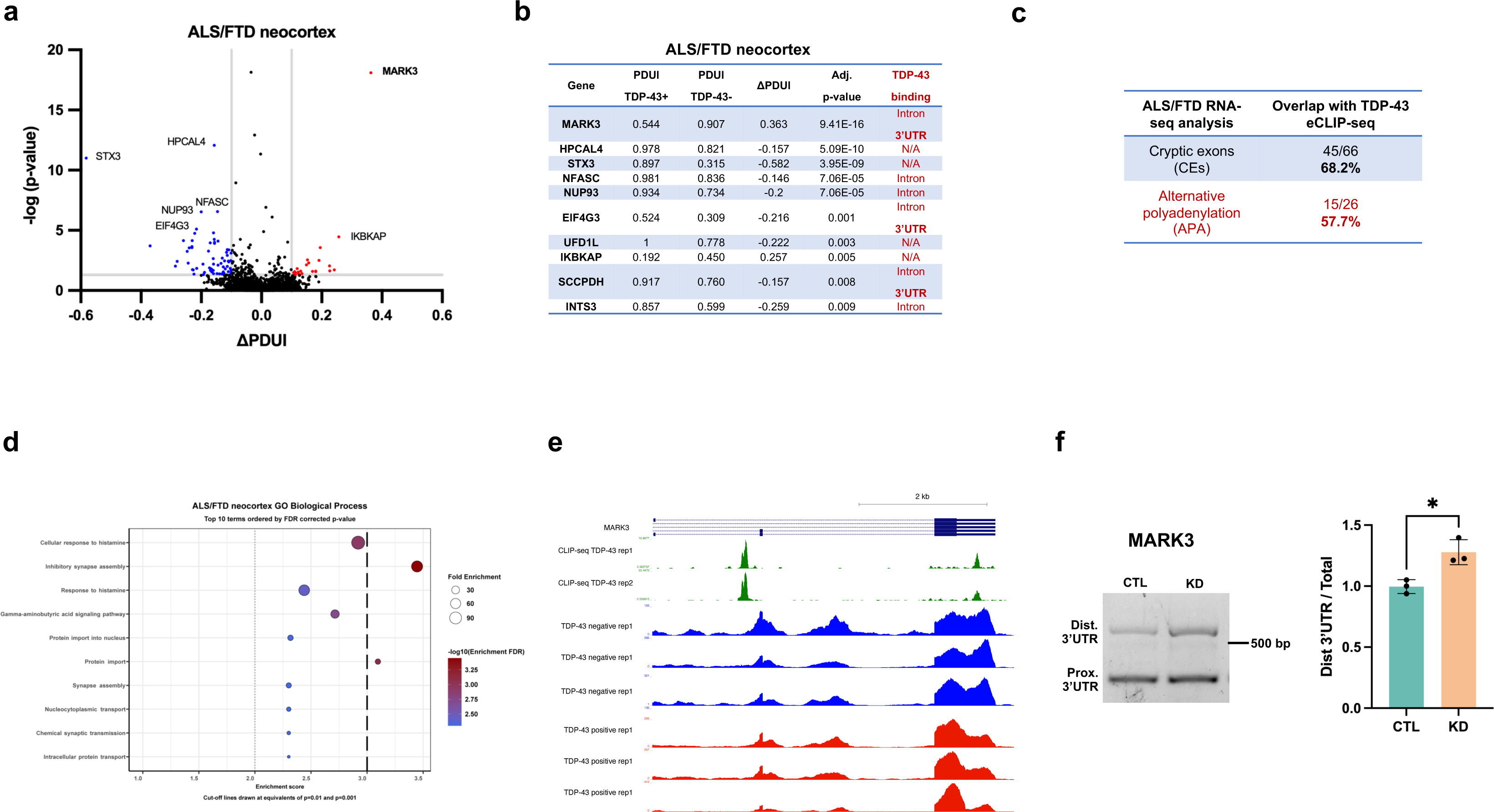
Nuclear clearance of TDP-43 induces APA in ALS/FTD patient neurons. (**a**) Volcano plot depicting APA genes in neuronal nuclei from 7 post-mortem ALS/FTD neocortex samples sorted by FACS for the presence or absence of nuclear TDP-43. APA genes with p < 0.05 and ΔPDUI ≤ -0.1 are depicted in blue and APA genes with p < 0.05 and ΔPDUI ≥ 0.1 are in red. Please note that *IKBKAP* is also known as *ELP1*, which is how this gene is referred to in the accompanying paper from Zeng et al. ^26^ (**b**) Top 10 APA events by FDR adj. p-value with a ƖΔPDUIƖ ≥ 0.1 along with evidence of TDP-43 binding sites by eCLIP-seq (red). (**c**) Percentage of cryptic exon (CE)-containing genes and APA genes that contain TDP-43 binding sites in ALS/FTD neuronal nuclei depleted of TDP-43. (**d**) GO analysis of biological processes (BP) enriched for coding APA events with p < 0.05 and ƖΔPDUIƖ ≥ 0.1 highlights genes that function in nucleocytoplasmic transport and synapse assembly. (**e**) ENCODE CLIP-seq data showing the location of TDP-43 binding within the MARK3 3’UTR, as well as in an upstream intronic region (green). MARK3 binding within the 3’UTR is immediately upstream of the distal shift in 3’UTR usage observed in TDP-43 negative neurons (blue vs. red RNA-seq tracks). (**f**) RT-PCR of distal 3’UTR (top band) and proximal 3’UTR (bottom band) of MARK3 in iPSC-MNs in which TDP-43 was knocked down with shRNA for 10 days. *p < 0.05; two-tailed t-test; n = 3 biological replicates and n = 3 technical replicates. Data are presented as mean values ± SEM.

The most substantial 3’UTR lengthening in ALS/FTD neuronal nuclei (ΔPDUI = +0.363), was observed in the gene encoding microtubule affinity regulating kinase 3 (*MARK3*) (**Fig. 5b**). Visualizing RNA-seq tracks from this experiment with overlayed eCLIP-seq generated by the ENCODE project ^33^, we confirmed that TDP-43 binds *MARK3* in its 3’UTR immediately upstream of a canonical ATTAAA polyA signal (hg38: chr14:103503803-103503809), as well as in an upstream intron (**Fig. 5e**). This suggests that nuclear TDP-43 normally represses use of a distal *MARK3* PAS, which then becomes preferentially utilized upon nuclear depletion of TDP-43. Importantly, significant 3’UTR lengthening in *MARK3* was also observed in our APA analysis in SH-SY5Y and SK-N-BE(2) cells (**Fig. 2c, d**), and was independently confirmed in an accompanying paper from Gitler and colleagues ^26^, indicating that increased utilization of a distal polyA site in *MARK*3 is a prominent effect in neuronal cells upon loss of TDP-43. Given the consistency of this result, we evaluated this phenotype in iPSC-MNs, and confirmed that TDP-43 knock-down can induce markedly increased use of a distal PAS in the *MARK3* gene (**Fig. 5e, f**).

### MARK3 APA yields increased mRNA stability and elevated protein levels

As MARK3 is a tau kinase associated with tau S262 phosphorylation in the early stages of Alzheimer’s disease pathogenesis ^23^, and because mutations in closely-related MARK4 can significantly increase AD risk, promote hyperphosphorylation of tau, and induce neuron toxicity ^34^, we sought to investigate the functional consequences of MARK3 APA in neurons. RT-PCR analysis revealed that TDP-43 knock-down results in increased use of the distal *MARK3* PAS in SH-SY5Y cells (**Fig. 6a**). Although steady-state expression of *MARK3* did not change upon TDP- 43 knock-down (**Fig. (6b**), we observed a significantly longer half-life for *MARK3* RNA upon TDP- 43 depletion in combination with actinomycin D treatment (**Fig. (6c**), indicating that use of a distal *MARK3* PAS affects transcript turnover. Importantly, increased RNA stability corresponded with elevated MARK3 protein levels in SH-SY5Y cells (**Fig. (6d**). To characterize the cis-regulatory elements within the *MARK3* 3’UTR that may affect its metabolism, we evaluated predicted miRNA binding sites using miRBD ^35^ and TargetScan ^36^, and we obtained predicted RBP binding motifs using RBPmap ^37^ (**Fig. (6e**). Both miRNA prediction tools highlighted a conserved binding motif for miR-142-3p within the *MARK3* 3’UTR (**Supplementary Table 8**). Intriguingly, this miRNA was found to be upregulated in ALS mouse models and in sporadic ALS patients; indeed, serum levels of miR-142-3p have been negatively correlated with ALS clinical outcomes ^38^. We also identified two conserved RBP binding motifs within the *MARK3* 3’UTR, which can only be utilized when polyadenylation occurs at a distal MARK3 PAS (**Fig. (6e, Supplementary** Fig. 6**, Supplementary Table 9**). The first motif is recognized by TDP-43, as well as by other RBPs (e.g. RBM24 and RBM38), while a second T/G-rich motif is predicted to be bound by several other RBPs, such as TIA1 and HuR (**Fig. (6e**). The presence of highly conserved miRNA and RBP binding motifs in the distal region of the *MARK3* 3’UTR suggests that differential recognition of short- versus long-3’UTR isoforms by cis-regulatory elements may account for observed changes in *MARK3* mRNA stability upon TDP-43 knock-down.

**Figure 6.**
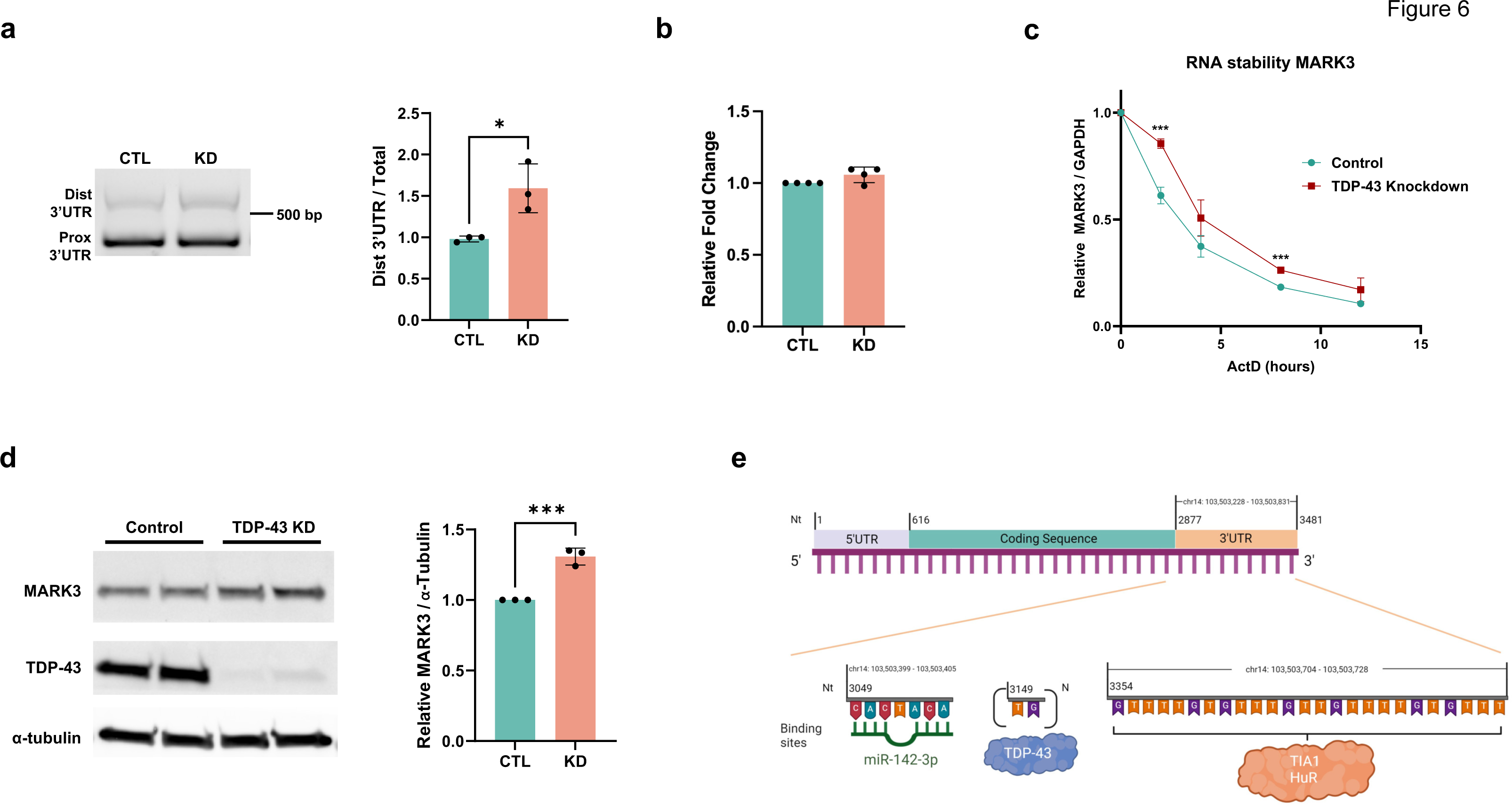
MARK3 alternative polyadenylation results in increased mRNA stability and protein levels in SH-SY5Y cells. **(a)** PCR validation of increased distal PAS usage of MARK3 upon TDP-43 KD in SH-SY5Y cells. **(b)** qRT-PCR of MARK3 in SH-SY5Y cells +/- TDP-43 knockdown. **(c)** RNA stability of MARK3 over time after actinomycin D (5 μg/mL) transcriptional arrest in SH- SY5Y cells upon TDP-43 depletion (red) or untreated (teal). **(d)** Immunoblot showing increased MARK3 protein levels upon TDP-43 KD in SH-SY5Y cells. **(a)** Graphical representation of select conserved cis-regulatory elements in the MARK3 3’UTR. *p<0.05, ***p<0.001; two-tailed t-test; n = 3 - 4 biological replicates. Data are presented as mean values ± SEM.

### Increased MARK3 expression promotes tau S262 hyperphosphorylation in iPSC-MNs

To determine if MARK3 protein levels are sufficient to drive tau S262 hyperphosphorylation in human motor neurons, we transduced iPSC-MNs with lentivirus encoding a shRNA vector against *MARK3*, lentivirus encoding the *MARK3* gene, or control empty vector lentivirus controls (**Supplementary** Fig. 7). In iPSC-MNs subjected to MARK3 over-expression, we detected a ∼2.4- fold increase in tau S262 phosphorylation (**Fig. 7a**). Moreover, we observed ∼50% reduction in tau S262 phosphorylation in iPSC-MNs subjected to MAP4K3 shRNA knock-down (**Fig. 7a**). When we performed TDP-43 shRNA knock-down in iPSC-MNs, we observed a trend towards increased MARK3 protein expression levels (**Fig. 7b**). Although knock-down of TDP-43 only elicited a trend toward increased MARK3 protein expression, we nonetheless measured a significant increase in tau S262 phosphorylation in iPSC-MNs subjected to TDP-43 shRNA treatment (**Fig. 7c**). These results reveal a potentially novel mechanistic link between TDP-43 and tau biology, and suggest that TDP-43 dysregulation of neuronal *MARK3* APA may contribute to altered cytoskeletal function in ALS/FTD and related neurodegenerative disorders.

**Figure 7:**
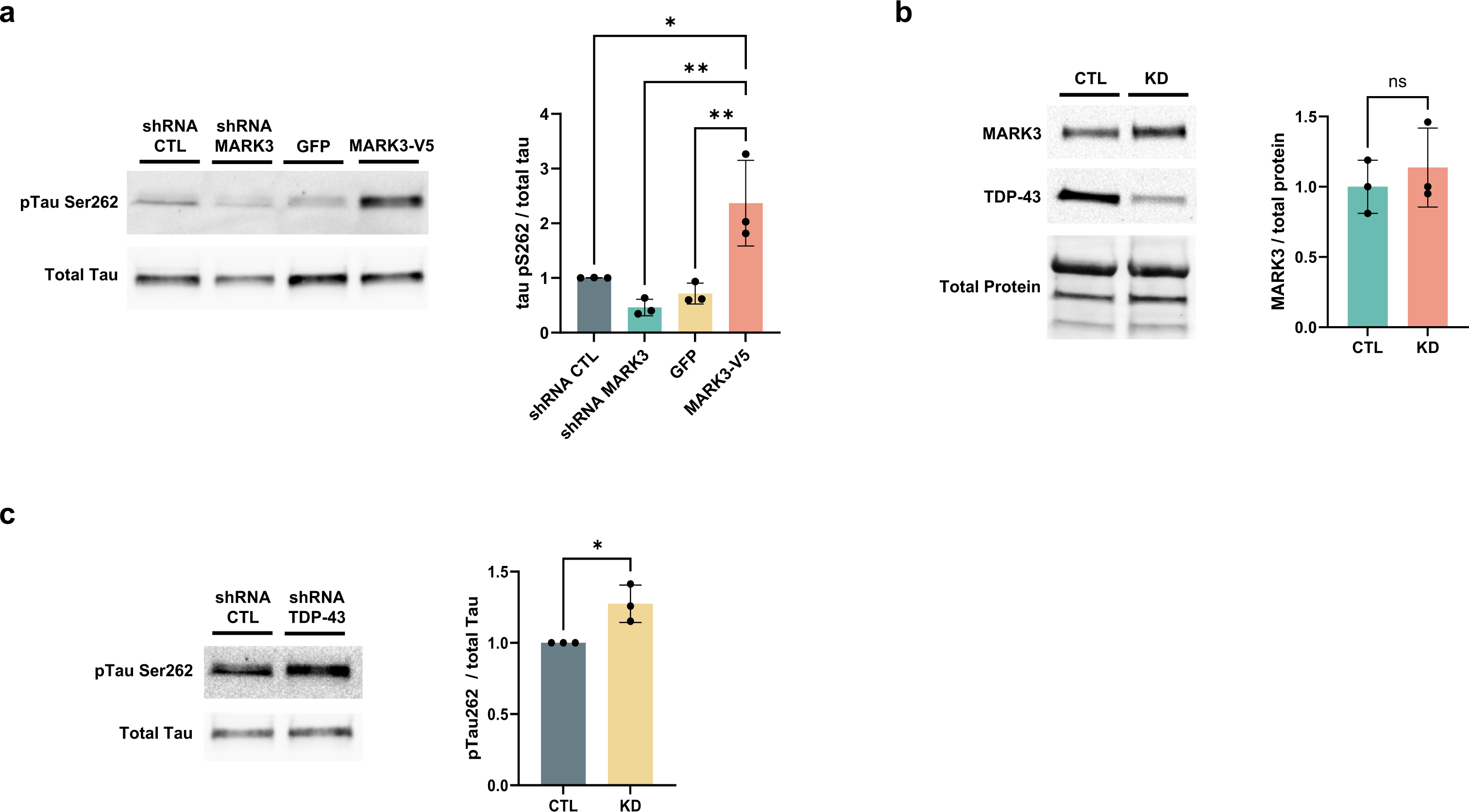
Increased MARK3 expression results in hyperphosphorylation of tau S262 in iPSC-MNs. (a) Immunoblot of S262 phosphorylated tau and total tau in iPSC-MNs transduced with lentivirus encoding control shRNA, shRNA MARK3, GFP (empty vector), or MARK3-V5 for 10 days. *p<0.05, **p<0.01; one-way ANOVA with post-hoc Tukey test; n = 3 biological replicates. Data are presented as mean values ± SEM. (**b**) Immunoblot of MARK3 in DIV 38 iPSC-MNs depleted of TDP-43 with shRNA for 10 days. *p<0.05, ***p<0.001; two-tailed t-test; n = 3 biological replicates. Data are presented as mean values ± SEM. (**c**) Immunoblot of S262 phosphorylated tau and total tau in iPSC-MNs transduced with lentivirus encoding control shRNA or shRNA TARDBP for 10 days. *p<0.05, ***p<0.001; two-tailed t-test; n = 3 biological replicates. Data are presented as mean values ± SEM.

## Discussion

The vast majority (>90%) of late-onset neurodegenerative diseases are sporadic, caused by ill-defined interactions between genetic and environmental risk factors. The complex etiology of these disorders has consequently hindered development of broadly effective therapies, and recent advances in the use of biological agents, which hold promise for treating rare familial forms of disease, may only benefit a small fraction of patients. This underscores the need to study convergent neurodegenerative disease mechanisms, such as the aberrant nuclear clearance and cytoplasmic aggregation of TDP-43, which has been implicated in a growing number of neurodegenerative disorders. Indeed, in addition to being a defining histopathological hallmark of ALS/FTD, emerging evidence suggests that TDP-43 dysfunction may play a pivotal role in dementia. Comorbid TDP-43 pathology is correlated with more severe cognitive impairment, more rapid disease progression, and increased brain atrophy in AD ^2,3^. Furthermore, TDP-43 has been shown to colocalize with senile plaques and neurofibrillary tangles, and TDP-43 histopathology is strongly associated with hippocampal sclerosis ^3^. Hence, delineating the cellular consequences of TDP-43 loss of function in neurons and other CNS cell types has the potential to be broadly relevant across multiple neurodegenerative diseases.

While the specific mechanisms linking TDP-43 dysfunction to neurodegeneration remain unclear, independent lines of investigation point towards aberrant RNA metabolism as the key driver of disease. Indeed, a number of independent studies have identified disease-relevant differentially expressed transcripts (e.g. *STMN2*) and alternatively spliced transcripts (e.g., *UNC13A*) that are being regulated by TDP-43 ^8,10–12^. TDP-43 is also known to regulate APA, and previous work in HEK293 cells defined a position-dependent principle by which binding of TDP-43 within 75 nucleotides of a PAS represses its usage, while binding further downstream of a PAS site enhances its usage ^13^. However, prior to this investigation and the two studies accompanying this report ^26,27^, the role of TDP-43 dysregulation of APA in neurodegenerative disease had not been undertaken. Here we found that TDP-43 loss of nuclear function changes polyadenylation site selection in hundreds of transcripts in neuron cell culture models of TDP-43 depletion or mutation and in postmortem ALS/FTD patient neuron nuclei containing or lacking TDP-43. While the majority of these APA events were cell-type specific, a number of highly reproducible changes in PAS usage were identified across multiple datasets, including significantly increased distal PAS usage in *CNPY3*, significantly reduced distal PAS usage in *SMC1a*, and significantly increased distal PAS usage in *MARK3*. Two reports accompanying this study also evaluated the effect of TDP-43 depletion in postmortem ALS/FTD patient neuron nuclei using an identical transcriptome data set ^28^, and despite employing distinct APA detection algorithms and bioinformatics analysis pipelines, both investigations identified hundreds of TDP-43 target genes subject to APA ^26,27^, underscoring the relevance of TDP-43 dysregulation of polyadenylation site selection to the disease process in TDP-43 proteinopathy. To assess the functional significance of TDP-43 APA in ALS/FTD disease pathogenesis, we also performed GO analysis on the APA hits and identified various pathways, including response to oxidative stress, which we validated as impaired in TDP- 43^N352S^ SH-SY5Y cells by documenting elevated ROS upon treatment with hydrogen peroxide.

To examine the functional implications of TDP-43 APA, we pursued directed studies on the tau kinase MARK3, our top hit from DaPars analysis of ALS/FTD patient post-mortem frontal cortex neuron nuclei containing or depleted of TDP-43. After confirming that TDP-43 knock-down in SH-SY5Y cells resulted in increased expression of a *MARK3* transcript isoform with a lengthened 3’UTR, we measured *MARK3* RNA expression levels, but did not detect any change in *MARK3* RNA levels upon TDP-43 knock-down. However, when we performed TDP-43 knock-down in combination with actinomycin D treatment to block transcription, we documented a significant increase in *MARK3* RNA expression levels, indicative of increased RNA transcript stability. Immunoblot analysis of MARK3 protein in SH-SY5Y cells subjected to TDP-43 knock-down revealed markedly increased expression of MARK3 protein, confirming that use of a distal PAS upon TDP-43 depletion results in increased MARK3 protein. Inclusions of tau protein accumulate in the brains of patients affected with familial and sporadic FTD ^39^, and tau is centrally involved in AD where as many as 57% of affected patients display TDP-43 nuclear clearance and cytoplasmic aggregation in addition to the hallmark neurofibrillary tangles of tau protein ^2^. To determine if TDP-43 APA of MARK3 could contribute to disease pathogenesis in ALS/FTD and AD, we evaluated the effect of TDP-43 knock-down on MARK3 phosphorylation of tau at S262, a well-established MARK3 phosphorylation site with potential pathophysiological implications, because tau phosphorylation at this residue, which is located within a tau microtubule-binding site ^40^, can reduce the microtubule-stabilizing properties of tau and antagonize microtubule tracking at the plus end by disrupting interaction between end-binding protein 1 and tubulin ^41^^42^. We found that TDP-43 knock-down in iPSC-derived human motor neurons resulted in significantly increased tau S262 phosphorylation, indicating that TDP-43 APA of *MARK3* could be contributing to neurodegeneration. Similarly, in an accompanying report from Zeng et al., TDP-43 APA of the FTLD-TDP risk gene *TMEM106B* was detected and then documented in post-mortem frontal cortex samples from FTLD-TDP patients, and proposed to alter *TMEM106B* translational efficiency, potentially destabilizing TMEM106B dimer formation ^26^.

Nearly two decades ago, the discovery of TDP-43 aggregates in the brains of patients with ALS and FTD revolutionized the neurodegenerative disease field ^43^, and set the stage for intense research into the role of TDP-43 dysfunction in the pathogenesis of ALS/FTD and related disorders. Over the last decade, considerable evidence has accumulated indicating that TDP-43 dysregulation of RNA splicing is likely a key step in the disease process for a broad range of disorders, now known as TDP-43 proteinopathies. However, in addition to regulating splicing, TDP-43 performs another fundamentally important function in the processing of RNA transcripts – selection of the site of polyadenylation, which dictates the length of a transcript’s 3’UTR. It is important to emphasize that 3’UTR length is a critical factor in regulating gene function. Within the 3’UTR are binding sites for miRNAs and RBPs, and these interactions dictate the stability of a transcript and hence modulate its expression at the protein level. Here we report that TDP-43 APA of MARK3 can alter its protein expression with consequences for tau function, underscoring the potential importance of TDP-43 dysregulation of polyadenylation site selection in disorders featuring tau pathology and TDP-43 nuclear clearance and cytoplasmic aggregation. Furthermore, cis-acting regulatory elements controlling mRNA subcellular localization exist within 3’ UTRs ^44^, and their presence or absence may determine if a mRNA localizes to the cell soma, the axon, or dendrite processes of a neuron. Subcellular localization of RNAs within neurons and other CNS cell types could profoundly affect cellular function; hence, altered subcellular mRNA transcript localization due to APA could be contributing to neurodegeneration, as reported for *ELK1* in an independent study from Bryce-Smith et al. accompanying this study ^27^. Hence, our results, along with independent findings accompanying this report from two studies using different methods of APA analysis ^26,27^, reveal that TDP-43 dysregulation of polyadenylation site selection could be driving the pathogenesis of various neurodegenerative diseases, likely in combination with altered splicing and cryptic exon inclusion. Defining genes subject to TDP-43 APA should thus be a major focus of future studies, as a number of these genes could be targets for developing biomarkers and engineering novel therapies. As antisense oligonucleotides (ASOs) can regulate polyadenylation site usage by steric hindrance ^45,46^, delineation of genes subject to APA may yield candidates for a next generation of ASOs and other biological agents for treatment of TDP-43 proteinopathy.

## Materials & Methods

### DaPars

software, described previously ^24^, allows for the joint analyses of multiple samples based on a two-normal mixture model, which is used to calculate the PDUI value. Briefly, we extract a 3′UTR annotation for each gene using the “DaPars_Extract_Anno.py” script within DaPars2. We then used the “samtools flagstat” command ^47^ to calculate the sequencing depth for each sample. Finally, we use DaPars2 to calculate the percentage of the distal poly (A) site usage index (PDUI) divided by the total expression level of each transcript across samples.

### Cell culture

Human neuroblastoma SH-SY5Y cells (ATCC) were maintained in 50% EMEM (ATCC, 30-2003), 50% Ham’s F12 (ThermoFisher, 11765047), supplemented with 10% fetal bovine serum, and 50 U/mL penicillin–streptomycin. Cells were grown at 37 °C and 5% CO2.

### Doxycycline-inducible knockdown of TDP-43

SH-SY5Y cells with stable integration of a doxycycline-inducible shRNA cassette targeting TDP- 43 were previously generated ^11^. Cells were treated with 1 μg/mL doxycycline (Sigma-Aldrich, D9891-1G) for 7 days to induce TDP-43 knockdown.

### iPSC-derived motor neurons (iPSC-MNs)

iPSCs from the KOLF2.1J line were maintained in mTeSR (STEMCELL Technologies, 100-0276). According to manufacturer’s instructions, iPSCs were differentiated into motor neurons from day 0-14 (STEMCELL Technologies, 100-0871) and subsequently matured from day 14- (STEMCELL Technologies, 100-0872). iPSC-MNs were transduced using lentiviral vectors at day 28 and collected at day 38. Cells were cultured at 37°C and 5% CO2.

### Lentiviral vector transduction

Lentiviral constructs encoding GFP (VectorBuilder, LVM[VB010000-9298rtf]-C ), MARK3-V5 (Addgene, 107235), shRNA control (Sigma, SHC002), tardbp shRNA (Sigma, TRCN0000016038), or MARK3 shRNA (Sigma, TRCN0000001564) were packaged into lentivirus by VectorBuilder. iPSC-MNs were transduced with 10 MOI lentivirus at DIV 28 for 10 days.

shRNA target sequences:

Control: CAACAAGATGAAGAGCACCAA Tardbp: GCTCTAATTCTGGTGCAGCAA MARK3: TGTGTGTGAAGTGGTGTATAT

### RT-PCR analysis

RNA was isolated from cell lysates (Qiagen, 74106) and reverse transcribed to generate cDNA (ThermoFisher, 11756500). PCR was performed using 10-30 ng cDNA template, 0.5 μM forward and reverse primers, and One*Taq* DNA polymerase mastermix (NEB, M0488L). PCR products were separated by agarose gel electrophoresis using 1.5% agarose (VWR, 0710) in 1X TBE buffer at 150V.

RT-PCR primer sequences:

MARK3 fwd: GGAAATGGAAGTGTGCAAGC

MARK3 prox rev: TAGAAGATGCAGACGTTATTGCC MARK3 dist rev: CACACACAAGCAATGTTCACAAC

CNPY3 var1 3’UTR fwd: CCAGCATCCTCTGTCCTGA CNPY3 var1 3’UTR rev: GAAGAGGGCACAGCCAAG CNPY3 var2 3’UTR fwd: GACCACCTGGGATCTTCCT CNPY3 var2 3’UTR rev: CACATCGTGGATCTTGCTGAG

SMC1A prox 3’UTR fwd: CCTGTCTGGATCCCTAAGCTG SMC1A prox 3’UTR rev: CTCCAGACCTAACATCACCTCTG SMC1A dist 3’UTR fwd: GTTAGTCAGTAGCAGTAGGAGGAG SMC1A dist 3’UTR rev: GCATTCACAGGGAAATAAGGAAGAC

### Quantitative reverse transcription PCR

Relative fold change of MARK3 was determined by qRT-PCR using SYBR Green Master Mix (ThermoFisher, A25776) with GAPDH as an endogenous control.

RT-qPCR primer sequences:

MARK3 fwd: CGCTCTCTGCTCCTCCTGTT MARK3 rev: TGCTGGTCTGACTCCTTTTCGG

GAPDH fwd: CAGTCTCCTCACCACAAAGTG GAPDH rev: CCATGGTGTCTGAGCGATGT

### Immunoblot analysis

Protein extracts were prepared in radioimmunoprecipitation assay lysis buffer (ThermoFisher, 89900) with protease and phosphatase inhibitors (ThermoFisher, 87786). Lysates were sonicated at 4 °C (Bioruptor Pico), denatured in LDS sample buffer (ThermoFisher, NP0007) with a reducing agent (ThermoFisher, NP0009), and heated at 70 °C for 10 min. Lysates were electrophoresed by sodium dodecyl sulfate–polyacrylamide gel electrophoresis using a 4%–20% gradient polyacrylamide gel (Bio-Rad, 5678093) and transferred to 0.45-μm polyvinylidene difluoride membrane (Bio-Rad, 1704157). Blots were incubated with primary antibodies diluted in 5% nonfat milk in TBST (TRIS-buffered saline, 0.05% Tween-20) overnight at 4 °C and with secondary antibodies diluted in 5% nonfat milk in TBST (ThermoFisher, A16078 [goat anti-mouse HRP, 1:3000], A16110 [goat anti-rabbit HRP, 1:3000], A32728 [goat anti-mouse Alexa FluorTM 647, 1:1000], A32733 [goat anti-rabbit Alexa FluorTM 647, 1:1000]) for 60 min at RT. Detection was performed using enhanced chemiluminescence substrate (Genesee Scientific, 20– 300S) and imaged on a ChemiDoc MP (Bio-Rad). Primary antibodies were as follows: TDP-43 (Proteintech, 12892-1-AP, 1:3000), MARK3 (Cell Signaling Technology, 9311S, 1:2000), SMC1A (abcam, ab9262, 1:2500), phospho-tau Ser262 (ThermoFisher, 44-750-G, 1:2000), TAU-5 (ThermoFisher, AHB0042, 1:2000), and α-tubulin (ThermoFisher, 62204, 1:3000 or Novus Biologicals, NBP2-80570, 1:3000).

### Actinomycin D mRNA stability experiment

SH-SY5Y cells with doxycycline-inducible TDP-43 shRNA expression were treated with 5 μg/mL actinomycin D (Sigma-Aldrich, A1410) for 0, 2, 4, 8, or 12 hours. Cells were subjected to a PBS wash prior to RNA isolation (Qiagen, 74106).

### Detection of reactive oxygen species (ROS)

Wildtype and TDP-43^N352S^ SH-SY5Y cells were plated on a 96-well plate and treated for 120 minutes with 100 μM H2O2. A fluorometric intracellular ROS detection kit was used to detect ROS levels at 30 and 120 minutes (Sigma, MAK144) using a Varioskan microplate reader (ThermoFisher).

### Gene Ontology (GO) analysis

GO analysis was performed using ShinyGO v0.77 ^48^ for GO Biological Processes and GO Molecular Function pathway databases. GO analysis was performed for APA events in protein-coding genes with ΔPDUI ≥ 0.1 and p < 0.05. Graphical representations of the top 10 pathways with an FDR adj. p <0.05 cut-off were generated in R version 4.3.1, utilizing the ggplot and sjPlot packages.

### Statistical analysis

As indicated, for pairwise comparisons, statistical significance was determined by two-tailed Student’s t-test. For multiple comparisons, statistical significance was determined by one-way analysis of variance (ANOVA) with *post hoc* Tukey test. The significance level (α) was set at 0.05 for all experiments. Data were compiled and analyzed using Microsoft Excel or Prism 10 (GraphPad).

## ACKNOWLEDGEMENTS

We wish to thank L. Stroud for excellent technical assistance, Dr. Don Cleveland (UC San Diego) and Dr. Ze’ev Melamed (The Hebrew University of Jerusalem) for kindly providing the TDP- 43^N352S^ homozygous SH-SY5Y cells, and Dr. Pietro Fratta (University College London) who generously provided the SH-SY5Y cells stably integrated with a doxycycline-inducible shRNA cassette targeting TDP-43. This work was supported by grants from the N.I.H. (R35 NS122140 to A.R.L.S., T32 AG000096 to F.J.A., R01 CA193466 to W.L., and RF1 NS118570 to M.G.H.), Chan-Zuckerberg Initiative Neurodegeneration Challenge Network (2021-239068 to A.R.L.S., W.L., and M.G.H.), Robert Packard Center for ALS Research at Johns Hopkins (PG12747 to A.R.L.S.), Muscular Dystrophy Association (Basic Science Development Award #865871 to F.J.A.), American Academy of Neurology (Richard Olney Clinician Scientist Development Award #23- CSDA-618 to F.J.A.), Medical Faculty of Ulm University (Clinician Scientist Program to S.M.), American Heart Association (Postdoctoral Fellowship #906383 to Y.C.), and Target ALS (Springboard Fellowship #FS-2023-SBF-S3 to F.J.A.).

## Author Contributions

F.J.A., Y.C., S.M. and A.R.L.S. provided the conceptual framework for the study. F.J.A., Y.C., S.M. O.H.T., M.G.H., W.L. and A.R.L.S. designed the experiments. F.J.A., Y.C., S.M., M.R.C., C.S., W.Y., S.M.,

W.G.S., K.C.K.E., and S.H. performed the experiments. F.J.A., Y.C., S.M. O.H.T., M.G.H., W.L. and

A.R.L.S. analyzed the data. F.J.A., Y.C., S.M. and A.R.L.S. wrote the manuscript.

## Declarations

M.G.H. is on the scientific advisory board of Transposon Therapeutics.

