## Supplemental Materials for "TDP-43 dysregulation of polyadenylation site selection is a defining feature of RNA misprocessing in ALS/FTD and related disorders"

### Supplementary Figure Legends

**Figure S1: APA events in SH-SY5Y cells and hESC-derived MNs upon TDP-43 knock-down**
(**a**) Table of top 10 APA events in SH-SY5Y cells treated with control siRNA or TARDBP siRNA for 96 hrs. (**b**) Depiction of ΔPDUI for all significant APA genes in TDP-43 depleted SH-SY5Y cells. (**c**) Overlapping APA genes between siRNA-treated SH-SY5Y cells and doxycycline-induced TARDBP knockdown SH-SY5Y cells, filtering for $p < 0.05$ and $|\Delta PDUI| \geq 0.1$. (**d**) Table of top 10 APA events in hESC-derived motor neurons (ESC-MNs) treated with control siRNA or TARDBP siRNA. Note that TARDBP siRNA treated ESC-MNs displayed only a partial (~60%) reduction in TDP-43 protein levels. (**e**) Depiction of ΔPDUI for all significant APA genes in TDP-43 KD ESC-MNs.

**Figure S2: Genes affected by APA function in ALS-relevant pathways**
Gene ontology (GO) analysis of biological processes (BP) and molecular functions (MF) of coding APA genes with $p < 0.05$ and $|\Delta PDUI| \geq 0.1$ in (**a, b**) SH-SY5Y cells, (**c, d**) SK-N-BE(2) cells, and (**e**) $i^3$Neurons (note that GO MF terms were not significantly enriched in $i^3$Neurons). Analysis was performed using ShinyGO v0.77.

**Figure S3: APA genes are bound by TDP-43 outside of the 3'UTR**
Venn diagram illustrating the intersection of APA events between SH-SY5Y cells, SK-N-BE(2) cells, and $i^3$Neurons depleted of TDP-43, along with evidence for direct TDP-43 binding in regions outside of the 3'UTR (5'UTR, intron, exon).

**Figure S4: RNA-sequencing tracks depict APA for the *CNPY3* and *SMC1A* genes in the presence or absence of TDP-43 knock-down**
RNA-seq tracks for *CNPY3* (**a**) or *SMC1A* (**b**) in $i^3$Neurons, SH-SY5Y cells, and SK-N-BE(2) cells in the presence or absence of TDP-43 knock-down. The location of the TDP-43 3'UTR binding site is depicted in green (*CNPY3* at chr6: 42,935,715 - 42,935,762; *SMC1A* at chrX: 53,379,722 - 53,379,785).

**Figure S5: Validation of TDP-43 knockdown in iPSC-MNs**
Western blot analysis of iPSC-MNs in which TDP-43 was knocked down with shRNA for 10 days.

**Figure S6: The MARK3 3'UTR contains two conserved RBP binding motifs**
Predicted RBP binding motifs in the MARK3 3'UTR with $p < 0.001$ and Z-score > 3.5 for at least one motif (RBPmap).

**Figure S7: Validation of MARK3 knockdown and overexpression in iPSC-MNs**

Immunoblot analysis of DIV 38 iPSC-MNs transduced with lentivirus encoding shRNA CTL, shRNA MARK3, GFP, or MARK3-V5 for 10 days. *$p < 0.05$, ***$p < 0.001$; two-tailed t-test; $n = 3$ technical replicates. Data are presented as mean values ± SEM.

**a**

| Gene | PDUI CTL | PDUI KD | ΔPDUI | p-value | Adj. p-value | TDP-43 binding |
|---|---|---|---|---|---|---|
| **ILF2** | 0.71 | 0.86 | 0.15 | 7.10E-10 | 1.22E-06 | N/A |
| **RPN2** | 0.883 | 1 | 0.117 | 8.77E-10 | 1.22E-06 | Intron |
| **HNRNPA1P10** | 0.75 | 0.86 | 0.11 | 1.26E-09 | 1.40E-06 | Exon |
| **SMC1A** | 0.28 | 0.0733 | -0.207 | 1.44E-06 | 0.000616 | Intron **3'UTR** |
| **PLD3** | 0.783 | 0.89 | 0.107 | 5.60E-05 | 0.0195 | N/A |
| **COASY** | 0.78 | 0.987 | 0.207 | 0.00012 | 0.0381 | N/A |
| **SSNA1** | 0.883 | 1 | 0.117 | 0.00015 | 0.0477 | N/A |
| **ACBD6** | 0.72 | 0.923 | 0.203 | 0.00024 | 0.0659 | Intron |
| **PNKD** | 0.817 | 0.987 | 0.17 | 0.00028 | 0.0754 | N/A |
| **STRA13** | 0.757 | 0.897 | 0.14 | 0.00045 | 0.104 | N/A |

**b**

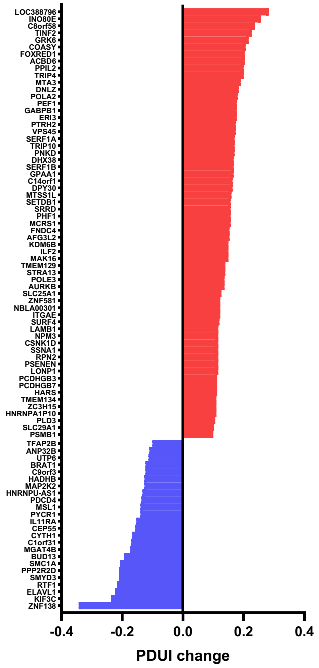


**c**

| Overlap with Brown et. al, 2022 |
|---|
| ACBD6 |
| ELAVL1 |
| GRK6 |
| INO80E |
| NPM3 |
| PPP2R2D |
| PTRH2 |
| SMC1A |
| ZNF138 |

**d**

| Gene | PDUI CTL | PDUI KD | ΔPDUI | p-value | Adj. p-value | TDP-43 binding |
|---|---|---|---|---|---|---|
| **SMC1A** | 0.722 | 0.467 | -0.255 | 0.0064 | 1 | Intron **3'UTR** |
| **GPCPD1** | 0.783 | 0.542 | -0.242 | 0.0055 | 1 | N/A |
| **PTPLB** | 0.907 | 0.692 | -0.215 | 0.011 | 1 | N/A |
| **CDK6** | 0.972 | 0.780 | -0.192 | 0.0053 | 1 | Intron Exon **3'UTR** |
| **DIDO1** | 0.515 | 0.327 | -0.188 | 0.043 | 1 | Intron |
| **AIG1** | 0.835 | 0.652 | -0.183 | 0.033 | 1 | Intron |
| **PLEKHM2** | 0.718 | 0.538 | -0.180 | 0.030 | 1 | N/A |
| **RANGRF** | 0.957 | 0.783 | -0.173 | 0.0009 | 0.55 | N/A |
| **TMEM11** | 0.937 | 0.765 | -0.172 | 0.0004 | 0.37 | N/A |
| **TFIP11** | 0.912 | 0.740 | -0.172 | 0.0093 | 1 | N/A |

**e**

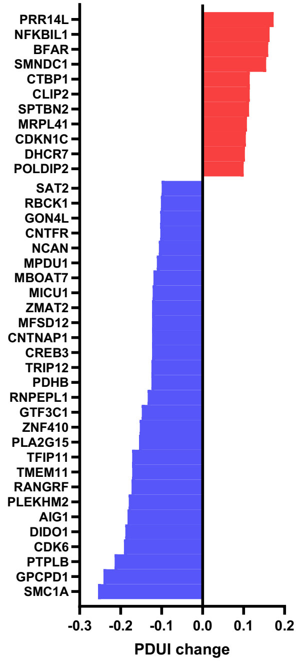


Figure S1

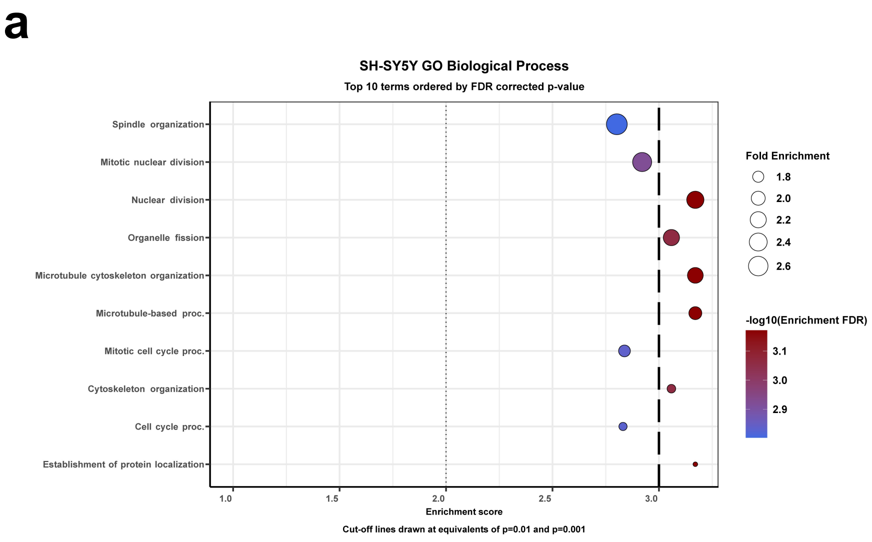
a
SH-SY5Y GO Biological Process
Top 10 terms ordered by FDR corrected p-value
Spindle organization
Mitotic nuclear division
Nuclear division
Organelle fission
Microtubule cytoskeleton organization
Microtubule-based proc.
Mitotic cell cycle proc.
Cytoskeleton organization
Cell cycle proc.
Establishment of protein localization
1.0
1.5
2.0
2.5
3.0
Enrichment score
Cut-off lines drawn at equivalents of p=0.01 and p=0.001
Fold Enrichment
1.8
2.0
2.2
2.4
2.6
-log10(Enrichment FDR)
3.1
3.0
2.9

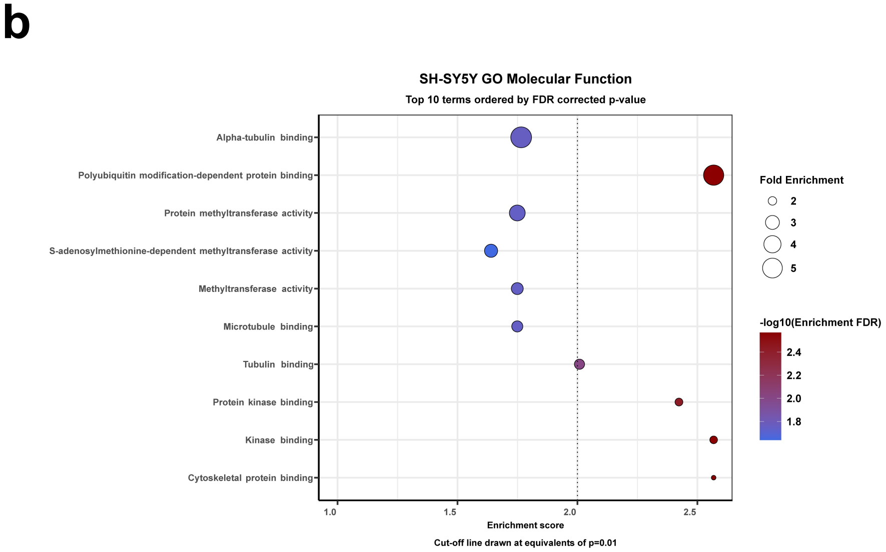
b
SH-SY5Y GO Molecular Function
Top 10 terms ordered by FDR corrected p-value
Alpha-tubulin binding
Polyubiquitin modification-dependent protein binding
Protein methyltransferase activity
S-adenosylmethionine-dependent methyltransferase activity
Methyltransferase activity
Microtubule binding
Tubulin binding
Protein kinase binding
Kinase binding
Cytoskeletal protein binding
1.0
1.5
2.0
2.5
Enrichment score
Cut-off line drawn at equivalents of p=0.01
Fold Enrichment
2
3
4
5
-log10(Enrichment FDR)
2.4
2.2
2.0
1.8

Figure S2

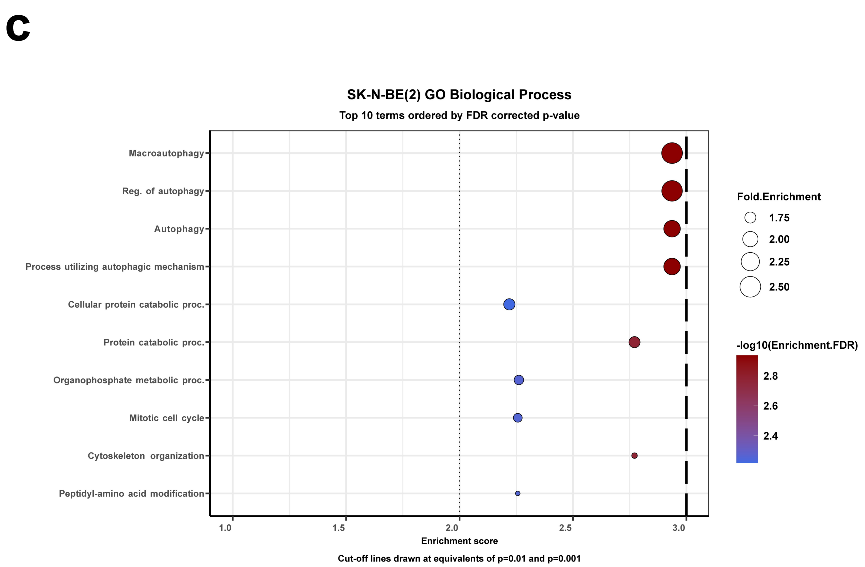
c
SK-N-BE(2) GO Biological Process
Top 10 terms ordered by FDR corrected p-value
Macroautophagy
Reg. of autophagy
Autophagy
Process utilizing autophagic mechanism
Cellular protein catabolic proc.
Protein catabolic proc.
Organophosphate metabolic proc.
Mitotic cell cycle
Cytoskeleton organization
Peptidyl-amino acid modification
1.0
1.5
2.0
2.5
3.0
Enrichment score
Cut-off lines drawn at equivalents of p=0.01 and p=0.001
Fold.Enrichment
1.75
2.00
2.25
2.50
-log10(Enrichment.FDR)
2.8
2.6
2.4

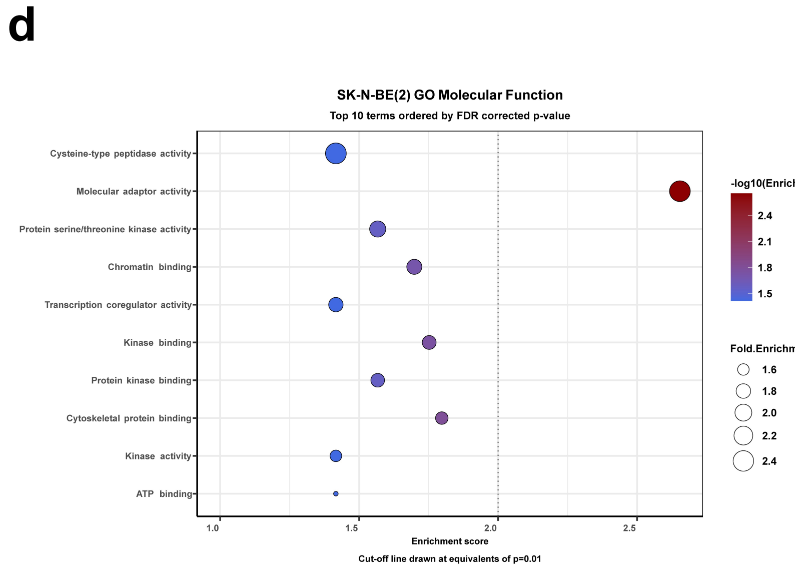
d
SK-N-BE(2) GO Molecular Function
Top 10 terms ordered by FDR corrected p-value
Cysteine-type peptidase activity
Molecular adaptor activity
Protein serine/threonine kinase activity
Chromatin binding
Transcription coregulator activity
Kinase binding
Protein kinase binding
Cytoskeletal protein binding
Kinase activity
ATP binding
1.0
1.5
2.0
2.5
Enrichment score
Cut-off line drawn at equivalents of p=0.01
-log10(Enricl
2.4
2.1
1.8
1.5
Fold.Enrichn
1.6
1.8
2.0
2.2
2.4

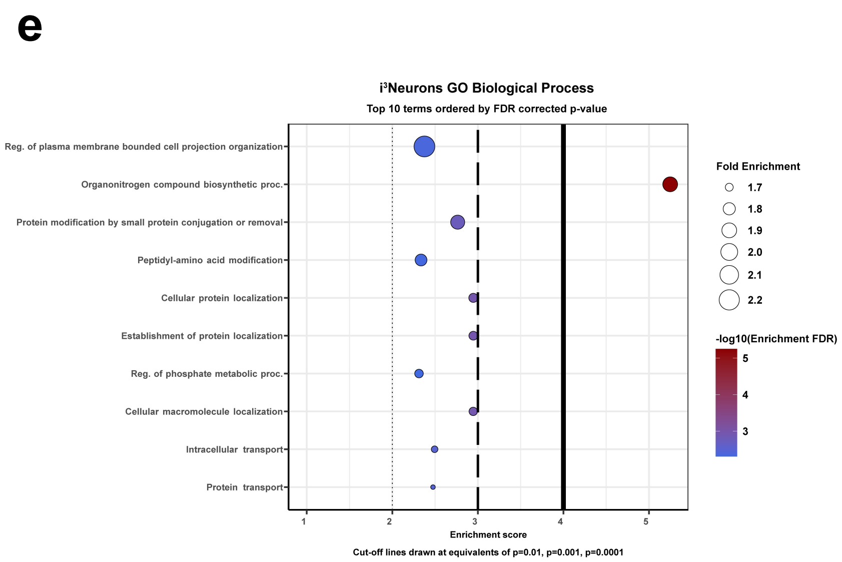
e
i³Neurons GO Biological Process
Top 10 terms ordered by FDR corrected p-value
Reg. of plasma membrane bounded cell projection organization
Organonitrogen compound biosynthetic proc.
Protein modification by small protein conjugation or removal
Peptidyl-amino acid modification
Cellular protein localization
Establishment of protein localization
Reg. of phosphate metabolic proc.
Cellular macromolecule localization
Intracellular transport
Protein transport
1
2
3
4
5
Enrichment score
Cut-off lines drawn at equivalents of p=0.01, p=0.001, p=0.0001
Fold Enrichment
1.7
1.8
1.9
2.0
2.1
2.2
-log10(Enrichment FDR)
5
4
3

Figure S3

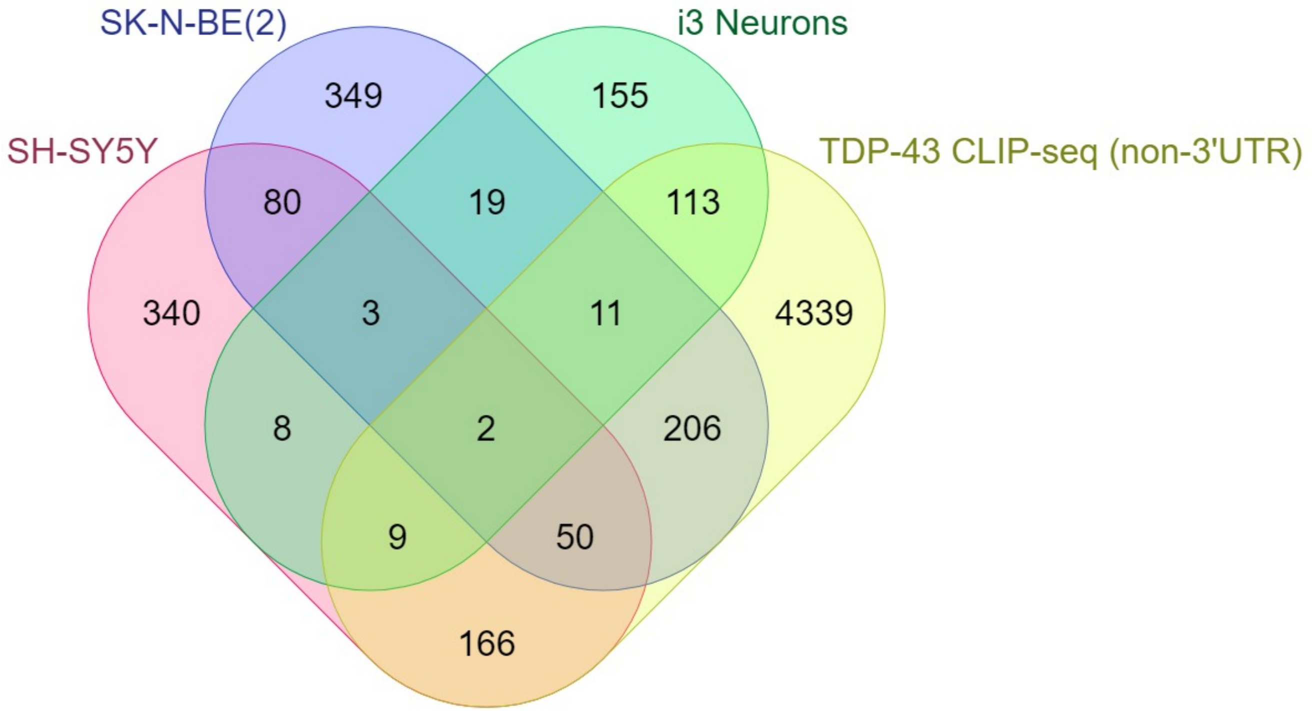
SK-N-BE(2)
i3 Neurons
SH-SY5Y
TDP-43 CLIP-seq (non-3'UTR)
349
155
80
19
113
340
3
11
4339
8
2
206
9
50
166

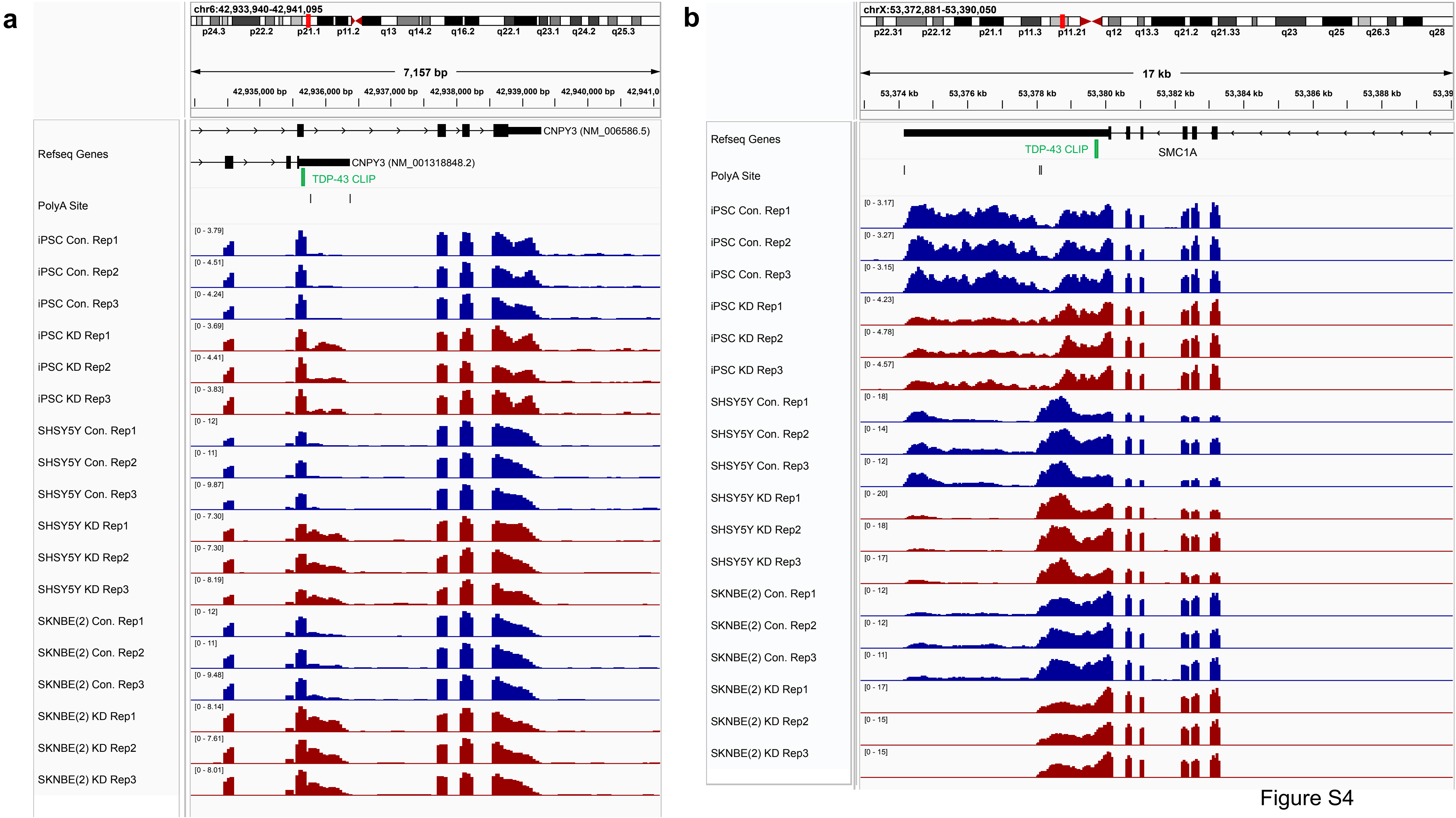
a
chr6:42,933,940-42,941,095
7,157 bp
Refseq Genes
CNPY3 (NM_006586.5)
CNPY3 (NM_001318848.2)
TDP-43 CLIP
PolyA Site
iPSC Con. Rep1
iPSC Con. Rep2
iPSC Con. Rep3
iPSC KD Rep1
iPSC KD Rep2
iPSC KD Rep3
SHSY5Y Con. Rep1
SHSY5Y Con. Rep2
SHSY5Y Con. Rep3
SHSY5Y KD Rep1
SHSY5Y KD Rep2
SHSY5Y KD Rep3
SKNBE(2) Con. Rep1
SKNBE(2) Con. Rep2
SKNBE(2) Con. Rep3
SKNBE(2) KD Rep1
SKNBE(2) KD Rep2
SKNBE(2) KD Rep3
b
chrX:53,372,881-53,390,050
17 kb
Refseq Genes
TDP-43 CLIP
SMC1A
PolyA Site
iPSC Con. Rep1
iPSC Con. Rep2
iPSC Con. Rep3
iPSC KD Rep1
iPSC KD Rep2
iPSC KD Rep3
SHSY5Y Con. Rep1
SHSY5Y Con. Rep2
SHSY5Y Con. Rep3
SHSY5Y KD Rep1
SHSY5Y KD Rep2
SHSY5Y KD Rep3
SKNBE(2) Con. Rep1
SKNBE(2) Con. Rep2
SKNBE(2) Con. Rep3
SKNBE(2) KD Rep1
SKNBE(2) KD Rep2
SKNBE(2) KD Rep3

Figure S4

Figure S5

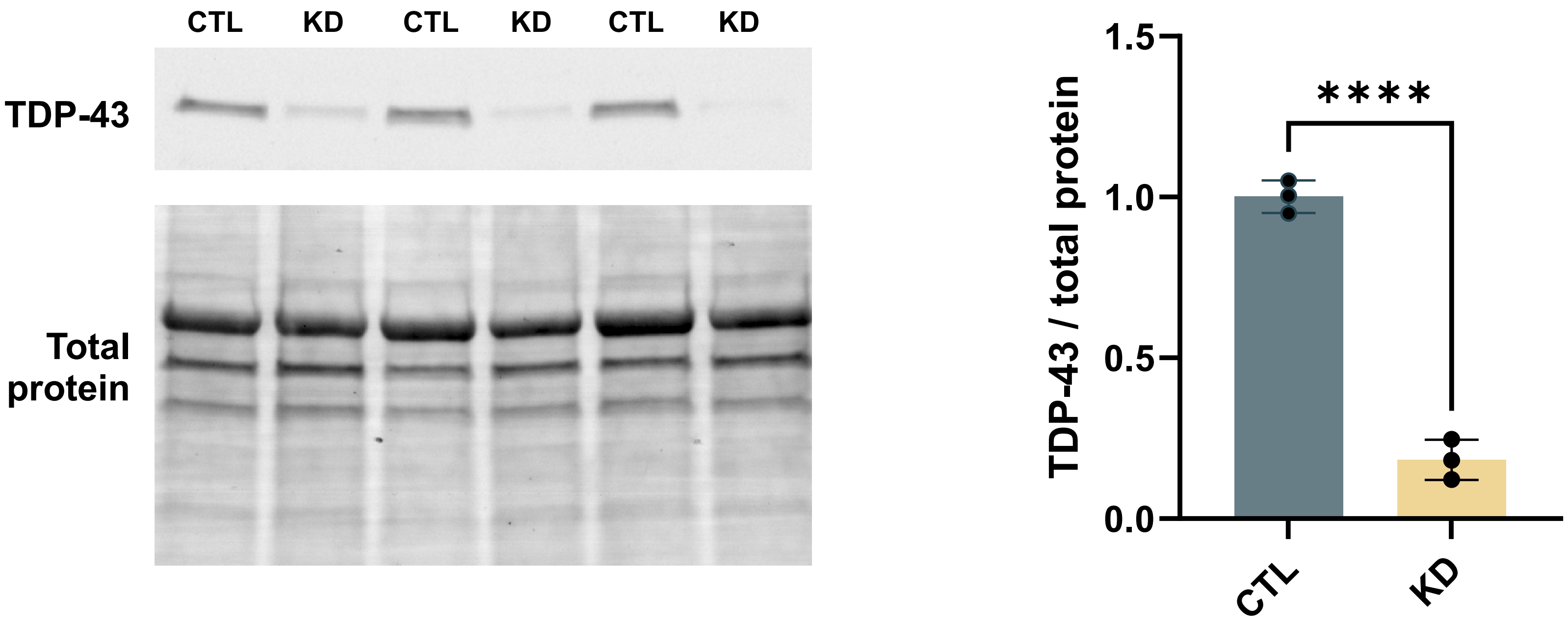
CTL
KD
CTL
KD
CTL
KD
TDP-43
Total protein
TDP-43 / total protein
1.5
1.0
0.5
0.0
****
CTL
KD

Figure S6

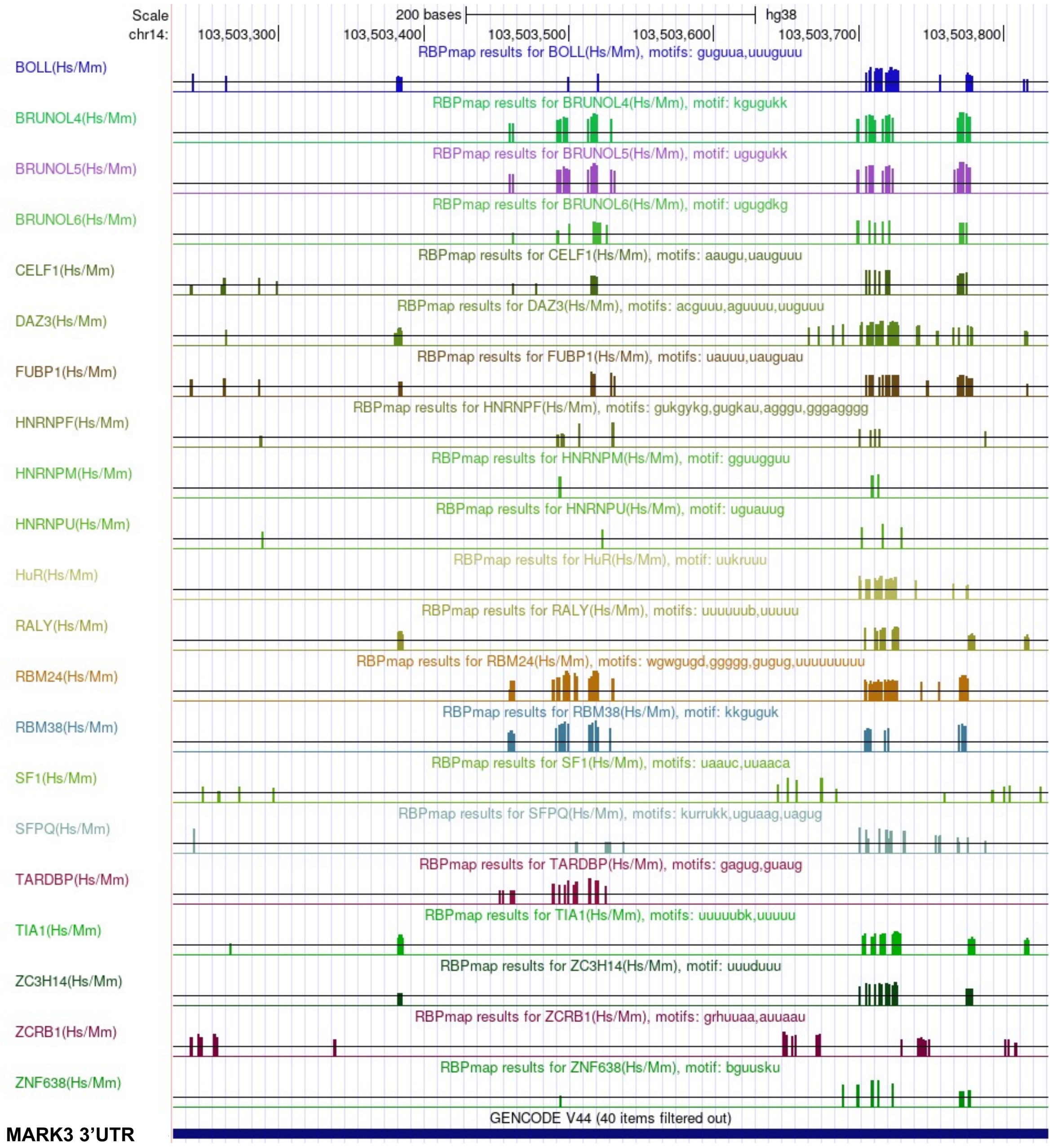
Scale
chr14:
200 bases
hg38
103,503,300
103,503,400
103,503,500
103,503,600
103,503,700
103,503,800
BOLL(Hs/Mm)
RBPmap results for BOLL(Hs/Mm), motifs: guguua,uuuguuu
BRUNOL4(Hs/Mm)
RBPmap results for BRUNOL4(Hs/Mm), motif: kgugukk
BRUNOL5(Hs/Mm)
RBPmap results for BRUNOL5(Hs/Mm), motif: ugugukk
BRUNOL6(Hs/Mm)
RBPmap results for BRUNOL6(Hs/Mm), motif: ugugdkg
CELF1(Hs/Mm)
RBPmap results for CELF1(Hs/Mm), motifs: aaugu,uauguuu
DAZ3(Hs/Mm)
RBPmap results for DAZ3(Hs/Mm), motifs: acguuu,aguuuu,uuguuu
FUBP1(Hs/Mm)
RBPmap results for FUBP1(Hs/Mm), motifs: uauuu,uauguau
HNRNPF(Hs/Mm)
RBPmap results for HNRNPF(Hs/Mm), motifs: gukgykg,gugkau,agggu,gggagggg
HNRNPM(Hs/Mm)
RBPmap results for HNRNPM(Hs/Mm), motif: gguugguu
HNRNPU(Hs/Mm)
RBPmap results for HNRNPU(Hs/Mm), motif: uguauug
HuR(Hs/Mm)
RBPmap results for HuR(Hs/Mm), motif: uukruuu
RALY(Hs/Mm)
RBPmap results for RALY(Hs/Mm), motifs: uuuuuub,uuuuu
RBM24(Hs/Mm)
RBPmap results for RBM24(Hs/Mm), motifs: wgwgugd,ggggg,gugug,uuuuuuuuu
RBM38(Hs/Mm)
RBPmap results for RBM38(Hs/Mm), motif: kkguguk
SF1(Hs/Mm)
RBPmap results for SF1(Hs/Mm), motifs: uaauc,uuaaca
SFPQ(Hs/Mm)
RBPmap results for SFPQ(Hs/Mm), motifs: kurrukk,uguaag,uagug
TARDBP(Hs/Mm)
RBPmap results for TARDBP(Hs/Mm), motifs: gagug,guaug
TIA1(Hs/Mm)
RBPmap results for TIA1(Hs/Mm), motifs: uuuuubk,uuuuu
ZC3H14(Hs/Mm)
RBPmap results for ZC3H14(Hs/Mm), motif: uuuduuu
ZCRB1(Hs/Mm)
RBPmap results for ZCRB1(Hs/Mm), motifs: grhuuaa,auuaau
ZNF638(Hs/Mm)
RBPmap results for ZNF638(Hs/Mm), motif: bguusku
GENCODE V44 (40 items filtered out)
MARK3 3'UTR

Figure S7

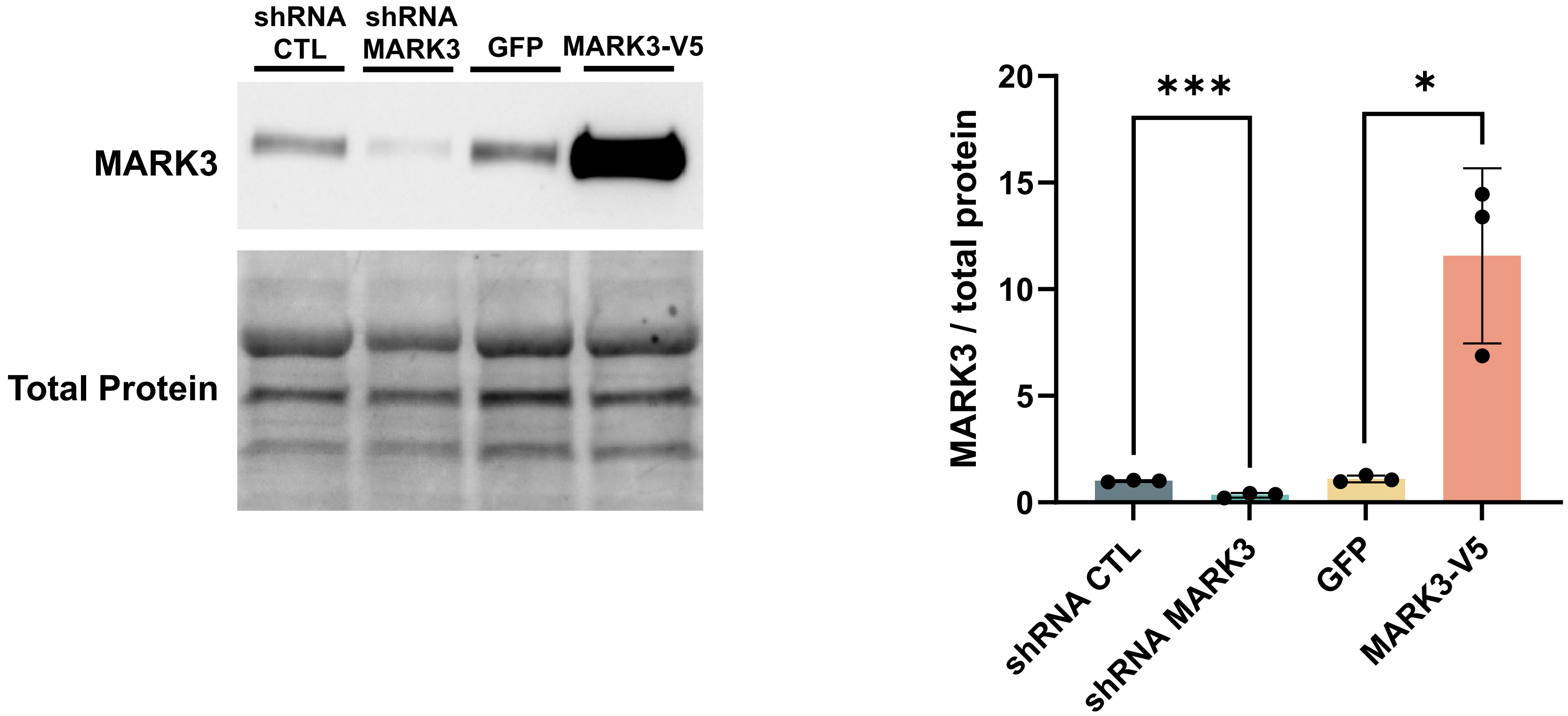
shRNA CTL
shRNA MARK3
GFP
MARK3-V5
MARK3
Total Protein
***
*
MARK3 / total protein
20
15
10
5
0
shRNA CTL
shRNA MARK3
GFP
MARK3-V5
